# Sensitive alignment using paralogous sequence variants improves long read mapping and variant calling in segmental duplications

**DOI:** 10.1101/2020.07.15.202929

**Authors:** Timofey Prodanov, Vikas Bansal

## Abstract

The ability to characterize repetitive regions of the human genome is limited by the read lengths of short-read sequencing technologies. Although long-read sequencing technologies such as Pacific Biosciences and Oxford Nanopore can potentially overcome this limitation, long segmental duplications with high sequence identity pose challenges for long-read mapping. We describe a probabilistic method, DuploMap, designed to improve the accuracy of long read mapping in segmental duplications. It analyzes reads mapped to segmental duplications using existing long-read aligners and leverages paralogous sequence variants (PSVs) – sequence differences between paralogous sequences – to distinguish between multiple alignment locations. On simulated datasets, Duplomap increased the percentage of correctly mapped reads with high confidence for multiple long-read aligners including Minimap2 (74.3% to 90.6%) and BLASR (82.9% to 90.7%) while maintaining high precision. Across multiple whole-genome long-read datasets, DuploMap aligned an additional 8-21% of the reads in segmental duplications with high confidence relative to Minimap2. Using Duplomap aligned PacBio CCS reads, an additional 8.9 Mbp of DNA sequence was mappable, variant calling achieved a higher F1-score and 14,713 additional variants supported by linked-read data were identified. Finally, we demonstrate that a significant fraction of PSVs in segmental duplications overlap with variants and adversely impact short-read variant calling.

## Introduction

High-throughput short-read sequencing technologies have transformed the study of genetic variation and the discovery of disease-associated variants for human disorders. However, the short read lengths (typically a few hundred bases) of short-read technologies such as Illumina limit the comprehensive detection of genetic variation [1]. The human genome is highly repetitive and contains several types of repetitive sequences including hundreds of long segmental duplications (ranging in length from a few kilobases to hundreds of kilobases) that have greater than 98% sequence similarity to other sequences [2, 3]. Some of these duplicated sequences are perfectly identical to their paralogous sequences over several kilobases. Duplications with length at least 10 kilobases and sequence identity of 98% or greater cover 3.0 *−* 3.2% of the human genome and overlap more than 800 protein-coding genes. Variants in many of these genes are implicated in rare Mendelian disorders as well as complex diseases [4]. Some examples of such duplicated genes are *PMS2* in Lynch syndrome [5], *STRC* in hearing loss [6], and *NCF1* [7] in autoimmune diseases. From the perspective of whole-genome sequencing, many of these segmental duplications are partially or completely inaccessible to short reads since the vast majority of reads originating from such regions cannot be unambiguously aligned to the genome [4, 8]. This limits the discovery of disease-associated mutations and our understanding of the function of these genes.

In recent years, two single molecule sequencing (SMS) technologies that can generate reads that are tens to hundreds of kilobases long have become widely available. The Pacific Biosciences (PacBio) SMRT technology can generate reads that are, on average, 10-60 kilobases long [9]. Another long read sequencing technology – Oxford Nanopore (ONT) MinION – can generate long reads with lengths that can even exceed a megabase in length [10]. The availability of these technologies has dramatically altered the ability to assemble bacterial and mammalian genomes since the long read lengths can resolve long repeats present in genomes [11]. The throughput and read lengths for these third-generation sequencing technologies continues to improve, as a result, these technologies are increasingly being used to sequence human genomes [12, 13]. The long read lengths of these technologies provide several advantages for sequencing human genomes compared to short reads. These include the ability to de novo assemble genomes with high contiguity [10, 14], reconstruct haplotypes directly from the sequence reads [15, 16] and increased sensitivity for the detection of structural variants [17, 12].

A key advantage of long SMS reads is their ability to map unambiguously in repetitive regions of the genome that include long segmental duplications with high sequence identity. This can enable accurate variant calling in these regions [16, 18]. However, variant calling using error-prone SMS reads is challenging and short-read variant calling tools do not work well for SMS reads [16, 18]. To address the challenge of variant calling using SMS reads with high error rates, several new methods [19, 18, 16, 20] have been developed. Some of these methods use deep learning based models [19, 20] to overcome the high error rate while others exploit the long-range haplotype information present in SMS reads to enable haplotype-resolved variant calling [16, 18]. Recent work has shown that these variant calling methods achieve high precision and recall for single nucleotide variant (SNVs) calling in unique regions of the human genome that is comparable to that using Illumina WGS [16]. More recently, Circular consensus sequencing (CCS) can generate long reads with high accuracy (99.8%) using multiple passes of the PacBio SMS technology over a single template molecule [14]. The high acccuracy of these HiFi reads enables the accurate detection of both SNVs and short indels in human genomes [14] and also improves the mappability of the genome (97.8% of non-gapped bases) compared to short reads (94.8%).

Nevertheless, many segmental duplications are much longer than the average read length of HiFi reads and remain difficult to map unambiguously [14]. Using simulated PacBio reads, Edge et al. [16] found that long-read alignment tools such as Minimap2 [21] and NGMLR [22] result in low recall for variant calling in segmental duplications. Long-read alignment tools typically calculate alignment or similarity scores for each of the possible mapping locations for a read and assign it a high mapping quality if the alignment score of the best location exceeds that of the second-best location using some threshold. Long repeated sequences in the human genome result in multiple locations with high scores and pose problems for long-read alignment tools. Recent work has shown that the accuracy of long-read mapping in extra-long tandem repeats in the human genome – typically found in centromeres – can be improved using specialized computational methods [23, 24, 25] that are designed to exploit the sequence and structure of long repeats. For example, the Winnowmap algorithm [24] modifies the sequence matching algorithm to avoid filtering out repeated *k*-mers that are common in tandem repeats [24].

In long segmental duplications with high sequence identity, there is potential to improve alignment accuracy by leveraging prior knowledge about the location and sequence of the duplications. Paralogous sequence variants (PSVs) – differences in sequence between a segmental duplication and its homologous sequences – are the primary source of information for assigning reads to their correct location in such regions. PSVs have previously been used to distinguish paralogous repeat copies and estimate paralog-specific copy number using short reads [26]. More recently, Vollger et al. [27] have developed a computational method for the de novo assembly of segmental duplications that uses PSVs to separate paralog copies. The high error rates of PacBio single-pass and ONT reads make the problem of distinguishing paralogous repeat copies even more difficult. In this paper, we describe a new probabilistic method for accurate mapping of long reads in segmental duplications that explicitly leverages PSVs to distinguish between repeat copies and assign reads with high confidence. Our method, DuploMap, builds on existing long-read alignment tools and carefully analyzes reads that are mapped to known segmental duplications in the genome. It performs local realignment around PSVs to maximally utilize the information present in noisy SMS reads.

PSVs are defined using a reference genome and it has been shown that a subset of PSVs correspond to polymorphisms in the human population [28, 29]. Using such unreliable or uninformative PSVs for differentiating repeat copies can result in conflicting evidence in support of different alignment locations resulting in reduced sensitivity and specificity of read mapping. To identify and discard uninformative PSVs, DuploMap jointly performs read mapping and PSV genotyping using an iterative algorithm. Only reliable PSVs are used to assign reads to homologous repeat copies. We use simulated data to evaluate the improvement in mappability using DuploMap on alignments generated using existing long-read mapping tools. We also demonstrate the impact of DuploMap on read mappability and variant calling in segmental duplication in the human genome using a number of real datasets generated using the Pacific Biosciences and Oxford Nanopore technologies. DuploMap is open-source software available at https://gitlab.com/tprodanov/duplomap.

## Materials and Methods

Given SMS reads aligned to a reference genome (using a long read aligner such as Minimap2), our objective is to analyze reads that overlap segmental duplications (and their homologous sequences), determine the most likely alignment location for each read and assign a mapping quality to it [31]. We assume that standard alignment tools can correctly align reads in the unique regions of the genome. Therefore, we do not examine reads that do not have a primary alignment overlapping segmental duplications. DuploMap uses prior knowledge about segmental duplications in the human genome to identify clusters of duplicated sequences and pairwise PSVs.

### Clustering segmental duplications and identifying PSVs

To identify segmental duplications and PSVs, we used a previously computed database of segmental duplications for the human genome [2, 3]. The database was downloaded from the UCSC table browser. First, we filtered out all pairs of homologous sequences for which the fraction of matching bases was less than 97% or the length of the alignment was less than 5000 bases. Next, we constructed a graph on the segmental duplications where each node was a genomic interval with homology to at least one other interval. This graph had two types of edges: (i) similarity edges between pairs of homologous sequences from the segmental duplication database and (ii) proximity edges between pairs of intervals that are less than 500 bases from each other. The proximity edges were added since reads that overlap intervals close to each other but not homologous have to be analyzed jointly since they affect the PSV genotypes of both components. For the hg38 reference human genome, the segmental duplication graph had 5,818 nodes and 26,301 edges (3,587 similarity and 22,714 proximity edges). We removed 88 of the 256 clusters that did not contain any duplications longer than 10 kilobases and with sequence similarity at least 98%. Most of the remaining clusters were small (less than 3 nodes) but the largest connected component contained 3,301 nodes and 17,752 edges.

To identify PSVs, we used Minimap2 (options -ax asm20) to align each pair of homologous sequences. For a pair of aligned sequences *S*_1_, *S*_2_, we first searched for anchors: *k*-mers shared between the homologous sequences and represented by *k* consecutive matches in the pair-wise alignment. Each anchor sequence is required toy be unique in a window around it (by default, *k* = 6 and window length = 20). This way we get a set of anchor starting positions 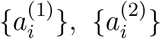. If the homologous sequences between two consecutive anchors are different: 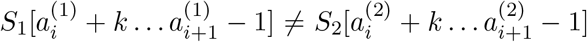, we define a pairwise PSV as a pair of intervals 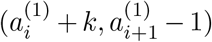 and 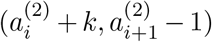. As a result of this, adjacent PSVs can be merged into a single PSV. The substrings 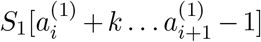 and 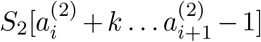 define the two alleles for the PSV. To avoid excessive number of PSVs in regions with low sequence similarity, we did not consider regions within the pairwise alignments that had sequence similarity lower than 95% and were longer than 300 bp. Finally, low-complexity PSVs (see Supplementary Methods for details) were filtered out and only high-complexity PSVs were retained for genotyping.

### DuploMap algorithm

For each cluster in the segmental duplication graph, DuploMap identifies reads that overlap segmental duplications in the cluster and analyzes the reads to determine the alignment location and mapping quality of each read. Unlike existing mapping tools, which map each read independently to the reference genome, DuploMap uses information from all reads jointly to align reads overlapping segmental duplications. This is done by identifying uninformative PSVs jointly with estimating the read alignment locations and mapping qualities. For a cluster of duplications, DuploMap first retrieves all reads for which the primary alignment intersect the genomic intervals contained in the cluster. Next, it performs the following steps on the set of reads:

1. For each read:

- Find the set of potential alignment locations,
- Use LCS-based filtering to discard some alignment locations,
- If the number of alignment locations after filtering is one, assign read to that location with high confidence (mapping quality = 254),
- Determine the actual alignment for the read and each alignment location using Minimap2.
2. For each PSV, estimate genotype likelihoods using reads aligned with high confidence (mapping quality greater than a threshold), and identify reliable PSVs.
3. For each read with two or more potential alignment locations, calculate location likelihoods using reliable PSVs and estimate mapping quality for best alignment location.
4. Repeat steps 2 and 3 until the read assignments do not change.

After we assign mapping locations and qualities for all reads for a given cluster of segmental duplications, we perform additional post-processing to identify reads that shown high rate of discordance with the genotypes of overlapping PSVs. Reads with high discordance can be result of missing duplicated sequences in the reference genome or due to other reasons such as structural variants. Reads that overlap at least five PSVs and show a high rate of discordance are assigned a low mapping quality (see Supplementary Methods for details). Next, we describe the individual steps (1, 2 and 3) of the algorithm in detail. The procedure for identifying the set of potential alignment locations uses the segmental duplication database (step 1) is described in the Supplementary Methods.

### Filtering alignment locations using longest common subsequences

For reads overlapping segmental duplications with high sequence identity, comparing the alignment scores for different candidate locations is not very informative of the correct location, particularly for reads with high error rates. We developed a LCS-based strategy that uses *k*-mers that are unique to a particular alignment location to filter out unlikely locations. The motivation underlying this approach is that the correct alignment location should share unique *k*-mers with the read, i.e. *k*-mers that are not present in other locations and the number of such shared unique *k*-mers should be significantly greater than other locations. This approach allows us to quickly map reads that have some part located outside segmental duplications as well as reads that intersect divergent region(s) within segmental duplications.

We use the LCSk++ algorithm [49] to find the longest common subsequence LCS_*k*_(*a, b*) of *k*-mers shared between a pair of sequences *a* and *b*. Function *N* (*·*) counts the number of non-overlapping *k*-mers in a set (see details in the Supplementary). Suppose, a read *r* has *n* candidate locations 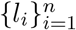. For a pair of locations *i* and *j* we find three LCS sets: LCS(*r, l*_*i*_), LCS(*r, l*_*j*_) and LCS(*l*_*i*_, *l*_*j*_). Let *A*_*ij*_ = *N* LCS(*r, l*_*i*_) *\* LCS(*r, l*_*j*_) be the *k*-mers that are present in the LCS between the read and the *i*-th location, but not in the LCS between the read and the *j*-th location. Additionally, let *B*_*ij*_ = *N (k*-mers(*l*_*i*_) *\* LCS(*l*_*i*_, *l*_*j*_) be the *k*-mers from the *i*-th location that are not in LCS(*l*_*i*_, *l*_*j*_). We use the Fisher’s Exact Test to calculate the *p*-value of the contingency table

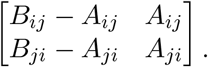

Without loss of generality, suppose that the read is more similar to the location *i* than to location *j*. In this case, a low *p*-value of the test would confirm that the ratio of shared read-location *k*-mers (*A*_*ij*_) to the number of unique *k*-mers for location *i* (*B*_*ij*_) is significantly higher than the corresponding ratio for location *j*: *A*_*ji*_/*B*_*ji*_. We considered values for *k* from 9 to 15, which showed similar results on simulated data (data not shown), and used *k* = 11. Since a single base difference can result in potentially *k* unique *k*-mers, counts of non-overlapping *k*-mers in the LCS are used for computing the Fisher’s exact test.

For a read with two or more alignment locations, we calculate the LCS-based *p*-value for each pair of locations. We say that location *i* dominates location *j* if *A*_*ij*_/*B*_*ij*_ > *A*_*ji*_/*B*_*ji*_ and the Fisher Exact Test *p*-value is less than a threshold (default = 0.0001). Then we select the smallest non-empty subset of locations that dominate all other locations using a directed graph (see supplementary methods).

### Read assignment using PSVs

For reads that have more than one possible alignment location after the LCS-based filtering, we use a PSV-based approach to determine the most likely alignment location (Figure 1b). For a long read *r* and *n* candidate alignment locations, we consider each pair of alignment locations in turn. For a pair of locations *i* and *j*, we use all high-complexity PSVs shared between these locations. For a PSV *v* we calculate read-PSV alignment probabilities for the two alleles of *v*. To account for the uncertainty in base-to-base alignment of long error prone reads, we use a small window around the PSV and average over all alignments using a pair-HMM [16]. We denote alignment probabilities 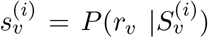 and 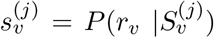, where *r*_*v*_ is read subsequence in a window around the PSV, 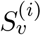 and 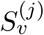 are the reference genome subsequences in the same window around the PSV at locations *i* and *j*. We cap alignment probabilities 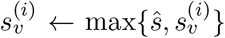 and 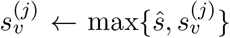 to reduce impact of a single PSV on read mapping, or a single read on a set of reliable PSVs 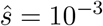 by default). Using these probabilities we calculate a likelihood for the true location of the read being *i* (relative to *j*):

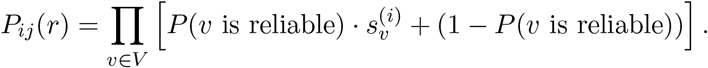

**Figure 1:**
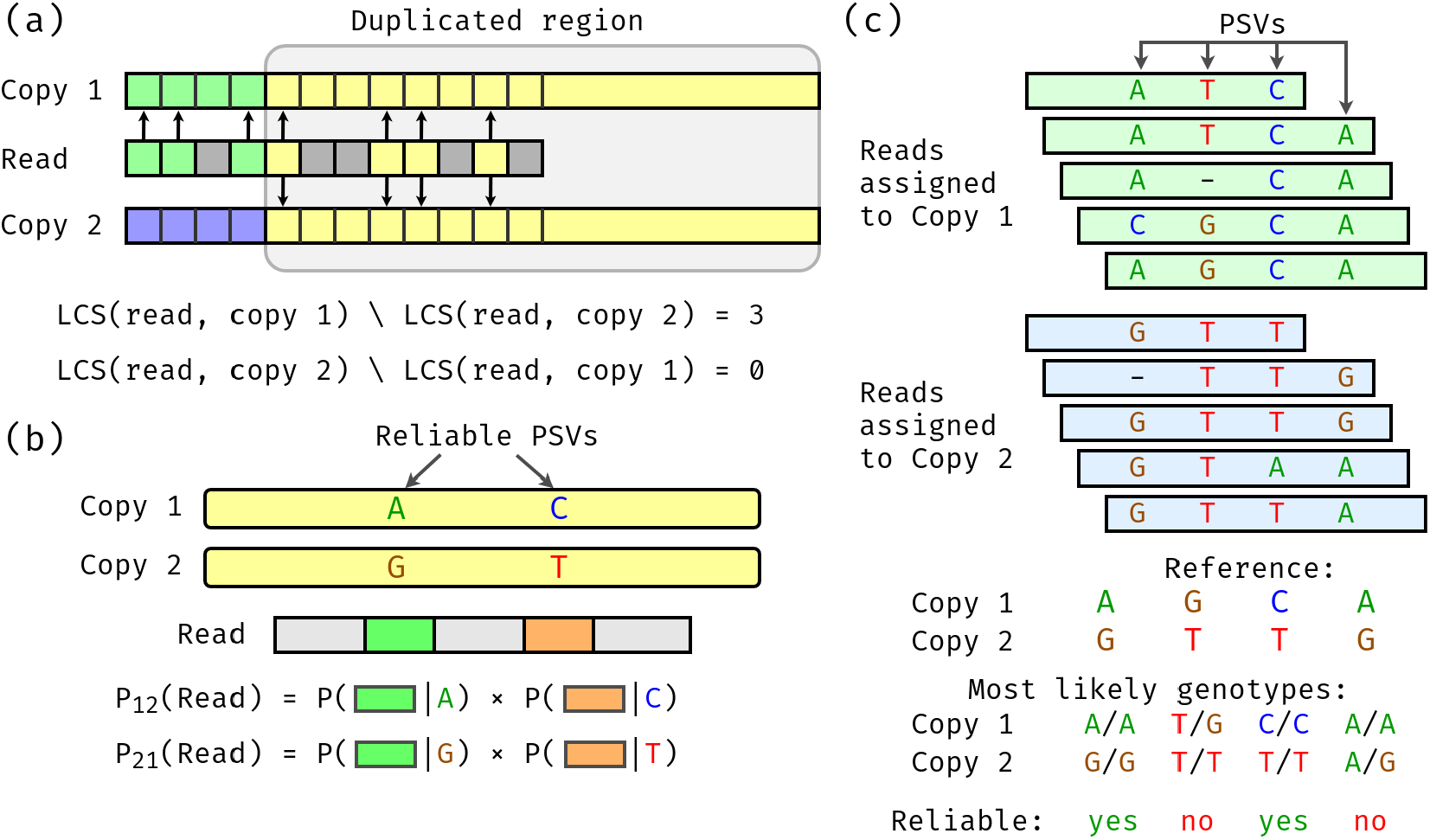
Overview of the DuploMap method. **(a) Filtering alignment locations using longest-common subsequences (LCS) of *k*-mers.** A read partially overlaps a segmental duplication and has two possible alignment locations (copy 1 and copy 2). The read and its possible locations are divided into *k*-mers that are shown with different colors. Arrows depict *k*-mers in the LCS between the read and the two copies. In the duplicated region, the read shares four *k*-mers with ‘copy 1’ that are also shared with ‘copy 2’. Outside the duplicated region, the read shares three *k*-mers (shown in green) with the *k*-mers of ‘copy 1’, but not with the *k*-mers of ‘copy 2’. **(b) Calculation of read-location probabilities using PSVs**. The read intersects two reliable PSVs that distinguish the two alignments locations. The probability of each location being correct (relative to the other location) are calculated using the local realignment probabilities between the read and the PSVs. **(c) Identifying reliable PSVs using assigned reads.** Five reads are mapped to ‘copy 1’ and five reads are mapped to ‘copy 2’ with high mapping quality. The genotype likelihoods for each PSV are calculated using these reads. Only two of the four PSVs have the reference genotype as the most likely genotype for both locations of each PSV and are considered reliable.

Note that *P* (*v* is reliable) is essentially the posterior probability of the reference genotype at both locations defined by a pairwise PSV. When the PSV is unreliable or uninformative, we assume that the PSV should be used to differentiate between the two locations and hence use a constant term (1) in the above equation. Initially, *P* (*v* is reliable) is assigned using a constant prior probability and in subsequent iterations it is estimated from the genotype likelihoods using reads assigned with high mapping quality to each location.

For reads with more than two candidate alignment locations, we use the pairwise likelihood to identify the “best” location *b* such that *P*_*bi*_(*r*) ≥ *P*_*ib*_(*r*) for all other alignment locations *i*. We select the second best location 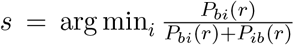 and assign the mapping quality as 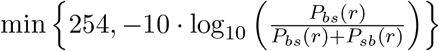. If no such location exists, we keep the original alignment of the read and assign it a mapping quality of 0.

### Identifying reliable PSVs using assigned reads

PSVs are defined using the reference genome sequence, however, since segmental duplications are difficult to assemble, some PSVs may be assembly artifacts. It is also possible that the analyzed genome has different alleles on homologous chromosomes for some of the PSVs, i.e. the PSV sites overlap with variants. For a PSV *v*, defined between two locations *i* and *j*, we select all reads *R*_*i*_ and *R*_*j*_ that cover the PSV and are assigned to location *i* and *j* respectively with high confidence (mapping quality greater or equal to a threshold). We use these reads to calculate the joint likelihoods of the genotype pair 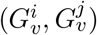 for the two locations. For each location, we consider three possible diploid genotypes defined by the two alleles of the PSV. For location *i* (*j*), the ‘0’ allele corresponds to the reference sequence of the PSV at location *i* (*j*) and the ‘1’ allele corresponds to the reference sequence of the PSV at location *j* (*i*). Hence the three possible genotypes for each location can be represented as *{0/0, 0/1, 1/1}* and we can estimate the posterior probability of each genotype pair (*g*_*i*_, *g*_*j*_) as follows:

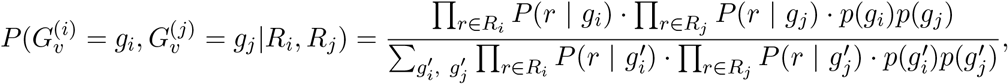

where *p*(*g*) is the prior probability of the genotype *g*. For most PSV sites, we expect the reference genotype (0*/*0) to be the correct one. Therefore, we assign a high value for *p*(0*/*0) and low probability for non-reference genotypes. For example *p*(0*/*0) = 0.95, *p*(0*/*1) = *p*(0*/*0) *·* (1 *−p*(0*/*0)) = 0.0475 and *p*(1*/*1) = 1 *−p*(0*/*0) *−p*(1*/*1) = 0.0025. Experiments on simulated data using values for *p*(0*/*0) ranging from 0.9 to 0.99 gave similar results (data not shown). Therefore, we use the prior value equal to 0.95 as default. *P* (*r | g*) is a probability of the read subsequence conditional on the genotype *g*, which is calculated using alignment probabilities:

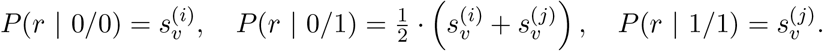

We define the probability *P* (*v* is reliable) as the posterior probability of the genotype being equal to the reference sequence at both locations, i.e. 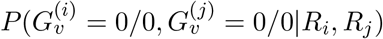.

### Measuring alignment accuracy

To assess the accuracy of read mapping in segmental duplications using simulated data, we used two metrics: (i) recall: the fraction of correctly mapped reads out of all simulated reads with true location overlapping *Long-SegDups*, and (ii) precision: the fraction of correctly mapped reads out of all reads mapped to *Long-SegDups*. A read is considered to be mapped to *Long-SegDups* if its primary alignment overlaps *Long-SegDups* with mapping quality greater or equal than a certain threshold. We say that a read is mapped correctly if it is mapped to *Long-SegDups*, its primary alignment covers the true location by at least 25% (this allows partial alignments, nevertheless the vast majority of the reads overlap the true location by more than 95% or less than 5%, see Supp. Figure 6). Additionally, the alignment should not go out of the true location by more than 100 bp in each direction to remove reads aligned to an incorrect copy in a tandem repeat. Precision and recall values for each mapping quality threshold were calculated by considering only reads with mapping quality greater than or equal to the threshold.

### Simulations

We used SimLoRD [30] to generate PacBio SMS reads (median lengths of 8.5 kb, 20 kb and 50 kb) from the reference human genome (hs37d5) using the default error rates of 0.11 for insertion, 0.04 for deletion, and 0.01 for substitution [30]. Reads were forced to only have a single sequencing pass to resemble PacBio CLR reads as opposed to CCS or HiFi reads. We aligned the SMS reads to the human reference (hs37d5) using the long-read alignment tools BLASR (`5.3.3`, options `--hitPolicy allbest --nproc 8`), MiniMap2 (`2.17-r941`, options `-t 8 -ax map-pb`) and NGMLR (`v0.2.7, options -t 8 -x pacbio`). To assess variant calling accuracy, we simulated a diploid genome using the reference human genome sequence with heterozygous SNVs (rate = 0.001) and homozygous SNVs (rate = 0.0005) [16].

### Whole-genome SMS datasets

We used whole-genome SMS datasets, generated using different sequencing technologies (PacBio CCS, PacBio CLR and Oxford Nanopore) for five individuals by the Genome in a Bottle (GIAB) consortium [33]. These datasets were downloaded from the GIAB ftp server: ftp://ftp-trace.ncbi.nlm.nih.gov/giab/ftp/data/ and were aligned to the hg38 reference genome using the tool Minimap2. In addition, Oxford Nanopore reads for NA12878 were obtained from the Nanopore WGS Consortium [10] and aligned to hg38 using minimap2. We also used 10X Genomics datasets (aligned reads and variant calls) for HG001 and HG002 obtained from the GIAB ftp server. Detailed information about the individual datasets is provided in Supplemental materials.

### Variant calling

Variants were called on the HG002 whole-genome PacBio CCS data using the tool Longshot [16] (v0.4.1). Variant calling was done using four different thresholds for the mapping quality (0, 10, 20 and 30). For each threshold, only reads with mapping quality greater than or equal to the threshold were used (-q parameter). Only variants with PASS filter and quality value at least 30 were used for analysis. High confidence variant call sets generated by the GIAB consortium were used for assessing accuracy of variant calling [33, 34]. For the HG002 genome, SNVs were compared against the GRCh38 version of the GIAB high-confidence call set (release v3.3.2 and v.4.1). The comparison of variant calls was limited to high-confidence regions (provided in a bed file). Precision and Recall were calculated using RTGtools vcfeval (v3.11). Comparison of different sets of variant calls was also done using RTGtools vcfeval.

## Results

### Overview of method

DuploMap is a probabilistic method specifically designed to improve the sensitivity and specificity of long-read alignments in segmental duplications in the genome. It starts from an existing set of aligned reads (generated using a long read alignment tool such as Minimap2) and updates the alignments and mapping qualities of reads that are mapped to segmental duplications. It utilizes a pre-computed database of segmental duplications and PSVs for this purpose. In the first step, DuploMap identifies the candidate alignment locations for each read whose initial alignment overlaps segmental duplications and uses an efficient filtering approach based on calculating the Longest Common Subsequence (LCS) to identify the most likely alignment location. This LCS-based filtering approach can identify the correct alignment location for reads that overlap a non-repetitive sequence (Figure 1a) and for reads from segmental duplications with moderate sequence identity. For reads that overlap segmental duplications with very high sequence identity, DuploMap aligns read sequences to possible alignment locations using Minimap2 and performs local realignment in the neighborhood of PSVs that overlap a read. The local realignments are used calculate read-location likelihoods and estimate the most likely location for each read (Figure 1b). Since some PSVs may not correspond to fixed differences between homologous sequences, DuploMap uses the reads assigned to each repeat copy (Figure 1c) to identify reliable PSVs – PSVs for which the genotype at the homologous positions is consistent with the reference genome. The read-location likelihoods are estimated using only reliable PSVs. Since reliable PSVs are not known in advance, the read assignments and the set of reliable PSVs are inferred using an iterative algorithm (see Methods).

Segmental duplications are defined as sequences with length at least 1 kb and sequence similarity *≥* 90% [3]. However, not all such segmental duplications are challenging for long read alignment. We used simulations to assess the mappability of SMS reads in duplicated sequences as a function of length and sequence similarity (data not shown). Based on these simulations, we constructed a subset of segmental duplications in the human genome with length greater than 5 kb and with sequence similarity at least 97%. These regions cover 86 and 101 megabases of the hg19 and hg38 reference human genomes, respectively. DuploMap only analyzes reads that overlap such segmental duplications (referred to as *Long-SegDups* in this paper).

### Evaluation of mapping accuracy using simulated reads

We simulated single-pass PacBio SMS reads using the SimLORD tool [30] with mean length equal to 8.5 kb and aligned them to the reference human genome using the long read alignment tool, Minimap2 [21]. Alignment tools report a mapping quality for each read which represents the probability that the reported alignment for a read is correct [31]: a mapping quality of 10 (20) corresponds to a probability of 0.9 (0.99). Analysis of the aligned reads showed that 74.9%, 69.0% and 63.1% of the reads that overlap *Long-SegDups* had mapping quality *≥* 10, *≥* 20 and *≥* 30 respectively. Furthermore, for reads completely within segmental duplications with *≥* 99.5% similarity, only 40.7% of reads had mapping quality greater or equal to 10. Although reads that overlap regions that are completely identical between the duplicated sequences cannot be mapped unambiguously, a significant fraction of the reads with low mapping quality overlapped multiple PSVs (illustrated for the *STRC* gene in Supp. Figure 1). Specifically, 70.6% of the reads that had a mapping quality less than 10 overlapped five or more PSVs. Next, we mapped the simulated reads using BLASR [32], a long read alignment tool developed specifically for PacBio reads. BLASR aligned a greater fraction of reads (80.8%) with mapping quality *≥* 20 compared to Minimap2 (68.4%) but was 28 times slower (Supplementary Table 1). This increased mappability came at the cost of accuracy: 3.2% of reads with mapping quality *≥* 20 were mapped to the incorrect location. The accuracy of another long read alignment tool, NGMLR [22], was significantly worse compared to Minimap2 and BLASR at all mapping quality thresholds (Supp. Figure 3).

Next, we used DuploMap to post-process the alignments generated using each of the long-read alignment tools separately. For a given mapping quality threshold, we used precision (fraction of correctly aligned reads out of reads *mapped* to *Long-SegDups*) and recall (fraction of correctly aligned reads out of *all simulated reads* in *Long-SegDups*) to assess the accuracy of read mapping. DuploMap improved both the precision and recall of read mapping in segmental duplications for all long-read mapping tools (Figure 2). For Minimap2, DuploMap improved the recall from 0.743 to 0.906, at a mapping quality threshold of 10, while maintaining high precision (0.9954, Figure 2 and Supp. Figure 2a). The improvement in recall was greater for higher mapping quality thresholds. Even if we consider all aligned reads (mapping quality threshold of 0), re-alignment using DuploMap increased both the precision and recall by 1.2 percentage points. DuploMap also improved both precision and recall for BLASR (Figure 2) and NGMLR (Supp Figure 3). In particular, the precision increased considerably from 0.965 (0.967) to 0.994 (0.995) while recall improved from 0.829 (0.806) to 0.907 (0.875) at a mapping quality threshold of 10 (20) for BLASR.

**Figure 2:**
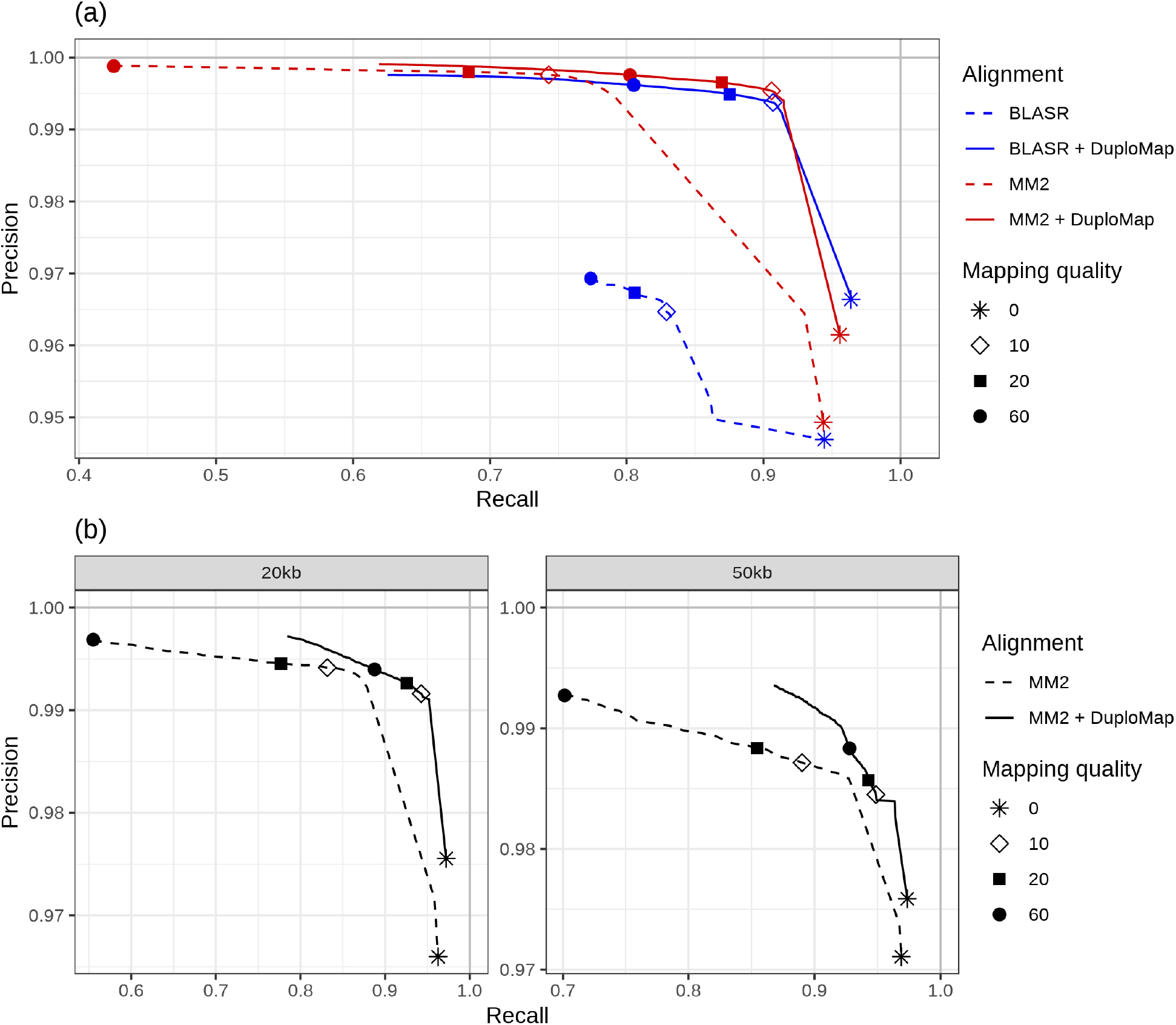
Accuracy of read mapping in segmental duplications using simulated long read data. Each curve shows the precision and recall of different alignment methods as a function of mapping quality thresholds. Dashed lines correspond to the original alignments while the solid lines show the alignments resulting from realignment using Duplomap. (a) Comparison of Minimap2 (MM2), BLASR, Minimap2+DuploMap, and BLASR+DuploMap on simulated reads with mean length 8.5 kb. (b)Accuracy of Minimap2 (MM2) and Minimap2+DuploMap on simulated reads with mean lengths 20kb and 50kb.

Minimap2 uses a minimizer based approach for finding matches between reads and the reference genome [21]. To improve speed, minimizers with a high frequency (top 0.02%) are discarded by default. We evaluated whether discarding a lower fraction (different values of the parameter *f*) could improve accuracy of read mapping in segmental duplications. Using *f* = 0 (use all minimizers) improved the recall slightly (0.743 to 0.763) but increased the memory usage five-fold (Supp. Figure 2b). Nevertheless, post-processing using DuploMap achieved a higher recall (0.906) while maintaining a high precision (0.9954). We also evaluated Winnowmap [24], a long-read alignment tool that uses a weighted sampling based method for selecting minimizers to improve long-read mapping using Minimap2 in long tandem repeats. However, Winnowmap’s recall and precision in *Long-SegDups* regions were lower than those for Minimap2 (Supp. Figure 3).

Next, we evaluated the accuracy of mapping SMS reads in segmental duplications as a function of read length. For this, we simulated PacBio single-pass reads of mean length 20 and 50 kilobases and aligned them to the reference genome using Minimap2. Not surprisingly, the recall for reads (at a fixed mapping quality threshold) increased as the read length increased (Figure 2b). Nevertheless, even for 50 kb long reads, recall was only 0.890 at a mapping quality threshold of 10. Re-alignment using DuploMap increased the recall to 0.949 for 50 kb reads, while keeping the precision high (0.985).

PSVs or paralogous sequence differences are defined using the reference genome sequence, however, some PSVs overlap with polymorphisms and should not be used to differentiate between the paralogous sequences. To assess whether DuploMap can map reads accurately in the presence of uninformative PSVs, we simulated PacBio reads from two-copy segmental duplications with 0%, 15% and 30% of the PSV genotypes assigned to be non-reference on one of the copies. DuploMap successfully identified uninformative PSVs and achieved high recall and precision for read mapping even with a high fraction of uninformative PSVs (Supp. Table 3).

### Improvement in read mapping for diverse SMS technologies

To assess the impact of DuploMap on read mapping in segmental duplications using real SMS data, we analyzed PacBio CCS whole-genome data for an individual, HG002 (NA24385), from the GIAB project [33]. The HiFi reads were initially aligned using Minimap2 to the hg38 reference genome. Post-processing the reads using DuploMap increased the percentage of reads with mapping quality *≥* 10 in segmental duplications from 65.7% to 80.6%, an increase of 15 percentage points (Figure 3). DuploMap can change both the alignment location and the mapping quality of reads overlapping segmental duplications. Comparison of the original Minimap2 and the DuploMap alignments showed that 4.8% of reads that had initially very low mapping quality (*<* 5) were aligned to a different location with mapping quality *≥* 30 (Figure 3). Similarly, DuploMap reduced the mapping quality of 1.9% of the reads – that initially had mapping quality *≥* 30 – to less than 10. We observed similar improvements in mappability for several human PacBio HiFi and CLR datasets (Table 1 and Supp. Figure 4). The increase in the percentage of reads aligned with a mapping quality greater than a threshold was consistently greater for HiFi reads compared to CLR reads. This was due to the improved ability to correctly allelotype PSVs using the HiFi reads: 1.7% local read-PSV alignments were ambiguous for CCS reads compared to 15.6% for CLR reads from the HG002 genome.

**Figure 3:**
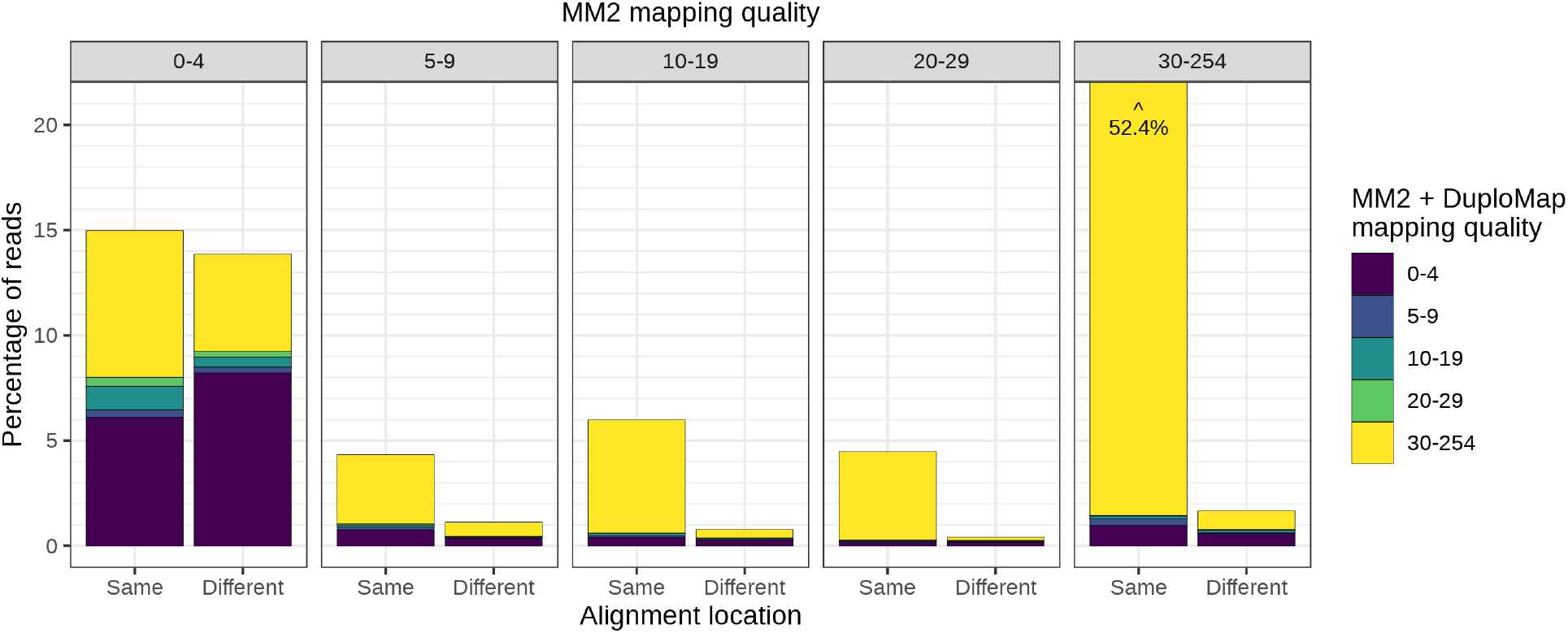
Comparison of mapping qualities and alignment locations for reads aligned with Minimap2 (MM2) and Minimap2+DuploMap on the HG002 CCS dataset. Five bar-lots corresponding to five bins of mapping quality using Minimap2 are shown. Each bar-plot shows the percentage of reads – color-coded by mapping quality after realignment using Duplomap – that had the same or different alignment location using Minimap2 and Minimap2+Duplomap. One of the bars (30-254 bin) that corresponds to 52.4% of reads is clipped for visual clarity.

**Table 1:** Improvement in mappability of reads using DuploMap on multiple SMS whole-genome sequence datasets. The last four columns show the percentage of reads with high mapping quality (*≥* 10 and *≥* 20) that overlap *Long-SegDups* regions in the *Minimap2* alignments and the difference between *Minimap2 + DuploMap* alignments and *Minimap2* alignments. CLR = Contiguous Long Reads, CCS = Circular Consensus Sequencing, ONT = Oxford Nanopore Technology, MM2= Minimap2.

| Genome | Sequencing technology | Median coverage | Reads analyzed | Read length (N50) | MM2 (%) | | $\Delta$ MM2+DuploMap (%) | |
| --- | --- | --- | --- | --- | --- | --- | --- | --- |
| | | | | | MQ $\geq 10$ | MQ $\geq 20$ | MQ $\geq 10$ | MQ $\geq 20$ |
| HG002 | PacBio CLR | 45 | 878k | 11,318 | 59.4 | 52.9 | +8.4 | +10.7 |
| HG003 | PacBio CLR | 20 | 416k | 10,999 | 59.9 | 53.5 | +9.8 | +11.3 |
| HG004 | PacBio CLR | 19 | 362k | 10,946 | 65.1 | 58.3 | +8.7 | +10.5 |
| HG002 | PacBio CCS | 29 | 300k | 13,480 | 65.7 | 58.9 | +14.9 | +19.5 |
| HG005 | PacBio CCS | 32 | 454k | 10,436 | 64.2 | 56.6 | +15.8 | +20.7 |
| HG001 | PacBio CCS | 29 | 381k | 10,004 | 71.6 | 63.7 | +15.0 | +21.2 |
| HG001 | ONT | 36 | 535k | 13,788 | 63.5 | 55.7 | +3.9 | +7.8 |
| HG002 | ONT | 58 | 464k | 54,352 | 64.5 | 58.0 | -1.5 | +1.7 |

Wenger et al. [14] demonstrated that relative to Illumina reads, PacBio HiFi reads increased the fraction of the genome that is mappable, i.e. covered by at least a certain number of reads with high mapping quality. Nevertheless, several disease-relevant genes such as SMN1 were still only partially mappable using HiFi reads. We assessed the impact of realignment using DuploMap on the mappable fraction of the human genome. To enable comparison between datasets with different sequencing coverages, we defined a genomic position as mappable if the number of reads covering it – with mapping quality greater than a threshold – is at least 50% of the median coverage for the dataset. Relative to Minimap2, realignment using DuploMap increased the fraction of the genome – limited to segmental duplications – that is mappable at all mapping quality thresholds (Figure 4). For HiFi reads, at a mapping quality threshold of 10 (20), 80.32% (79.47%) of the *Long-SegDups* regions were mappable relative to 69.01% (62.26%) using Minimap2. This also increased the mappability of 11 (16) of the 193 disease-associated duplicated genes using HiFi reads (Supp. Table 5).

**Figure 4:**
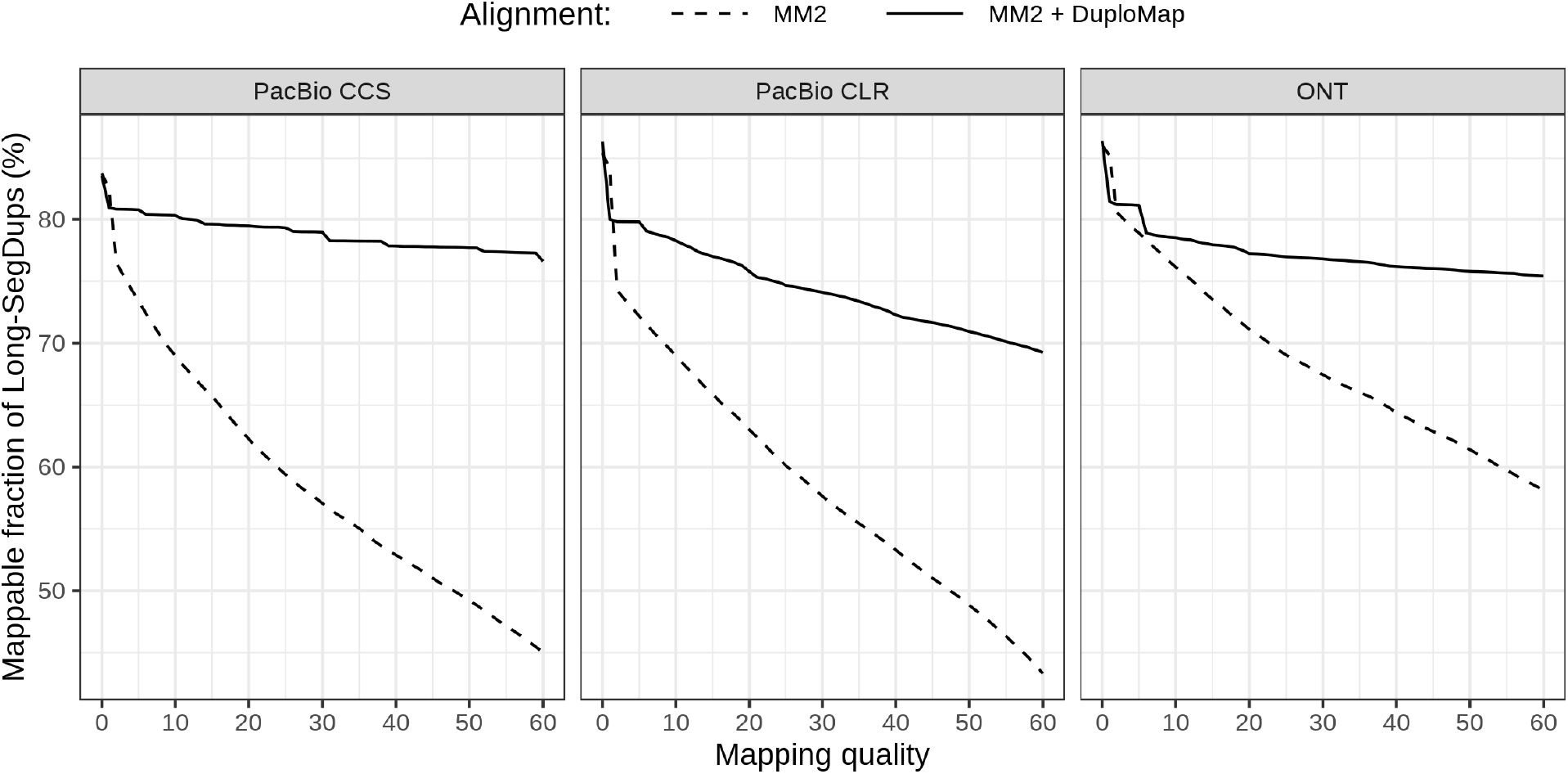
Improvement in mappability of *Long-SegDups* regions using DuploMap and three long-read datasets for the HG002 genome. Each sub-plot shows the percentage of the *Long-SegDups* regions (78.7 Mb on chromosomes 1-22) that is mappable at different mapping quality thresholds using *Minimap2* and *Minimap2 + Duplomap* alignments. A position is considered mappable if the number of reads covering it is at least 50% of the median coverage for the dataset.

Next, we analyzed a whole-genome human dataset generated using the Oxford Nanopore technology [10, 34]. Similar to PacBio datasets, realignment using DuploMap increased the fraction of reads with mapping quality greater than 10 (20) by 3.9 (7.8) percentage points (Table 1). For the ONT dataset with ultra-long reads (mean read length of 54.4 kb), only a minor improvement in the number of reads with high mapping quality was observed (Table 1). Nevertheless, at a mapping quality threshold of 10, an additional 1.9 Mb of DNA sequence is mappable using DuploMap aligned reads compared to reads aligned using Minimap2 (Figure 4).

DuploMap is multi-threaded and can use multiple cores to process clusters of segmental duplications in parallel. It required 2-5 hours (using 8 CPU cores) to process simulated and real whole-genome PacBio datasets with 30*×* coverage (Supp. Tables 1 and 2). This additional run-time was only 25-30% of the run-time of Minimap2 for generating the initial set of alignments. Since DuploMap infers reliable PSVs jointly using reads mapped to a cluster of segmental duplications, it needs to store all read alignments for a cluster in memory and hence the memory usage increases with increasing coverage (see Supp. Table 2).

### Variant calling in segmental duplications using DuploMap alignments

DuploMap increases the fraction of the genome - limited to segmental duplications - that is mappable using SMS reads. This is expected to improve the sensitivity of variant calling in such regions. To assess this, we used the variant calling tool Longshot [16] on simulated PacBio reads (30*×* coverage). SNVs were called using Longshot for four different mapping quality thresholds (0, 10, 20 and 30), i.e. reads with mapping quality below the threshold were not used for variant calling. We found that the recall for variants called in *Long-SegDups* using reads re-aligned with DuploMap was greater than that obtained using Minimap2-aligned reads at all mapping quality thresholds (Supp. Figure 5). At a mapping quality threshold of 10, the recall increased from 0.833 to 0.945 while the precision was virtually unchanged (*≥* 0.999 for both sets of alignments). The recall for Minimap2 was highest (0.898) when using all aligned reads (ignoring mapping quality) but resulted in significantly lower precision 0.904. For both Minimap2 and DuploMap, the best precision-recall tradeoff was observed at a mapping quality threshold of 10. For segmental duplications with 99.9% or greater identity, variants called using Minimap2+DuploMap alignments (mapping quality threshold of 10) had a recall 0.716 and precision equal to 0.998, compared to 0.253 and 0.994 respectively for variants obtained using Minimap2 alignments.

Next, we assessed the impact of the improved read mapping on variant calling using whole-genome PacBio HiFi data for HG002 (29*×* coverage). A recently developed variant calling tool, DeepVariant, has been shown to achieve very high precision and recall for PacBio HiFi reads [14]. Since a subset of the HG002 HiFi dataset was used for training the DeepVariant model [14], we used Longshot [16] for variant calling. Longshot has been shown to achieve high accuracy (F1 score of 0.9985) for SNV calling on CCS reads [35]. Across chromosomes 1-22, Longshot called 3,727,419 SNVs using the reads realigned with DuploMap (mapping quality threshold of 10), 18,291 more than using the Minimap2 aligned reads. We used the high-confidence benchmark variant calls from the GIAB consortium (v3.3.2) that cover approximately 2.35 Gb of the GRCh38 version of the reference genome (excluding the X and Y chromosomes) to assess the accuracy of the SNV calls. The precision and recall of SNV calling using the Minimap2-aligned reads and the DuploMap-aligned reads was identical: 0.9963 and 0.990 respectively (see Table 4). This was not surprising since the GIAB high-confidence benchmark variant calls (v3.3) were primarily generated using short read datasets and exclude the vast majority of repetitive regions in the genome.

The GIAB consortium recently released high-confidence benchmark variant calls (v4.1) for the HG002 genome that cover an additional 6% of the genome compared to the v3.3.2 calls. These benchmark variant calls incorporate information from 10X Genomics linked-read and PacBio HiFi read datasets and include variant calls in some segmental duplications. The precision and recall in the v4.1 regions for DuploMap and Minimap2 alignments were similar, although, SNV calls from DuploMap aligned reads had a higher F1 score (0.9905) compared to Minimap2 (Supp. Table 4). In the subset of the v4.1 regions that overlap *Long-SegDups* regions, DuploMap based calls had a higher recall compared to Minimap2 but lower precision at all mapping quality thresholds (Figure 5). Manual inspection of some of the false positives called using DuploMap aligned reads suggested that these may correspond to missing true positives in the GIAB v4.1 callset (see Supp. Figure 9). Hence, the true precision may be higher. Nevertheless, the F1 score of the DuploMap-based calls was consistently higher that the F1 score of the calls using Minimap2 alignments. In addition, the improvement in the F1 score was not dependent on the variant quality threshold used for Longshot (Supp. Table 4). Visual inspection of the SNVs calls that were called only using the DuploMap alignments and matched the v4.1 benchmark calls showed that the vast majority of these SNVs were not called using Minimap2 alignments due to low mapping quality of the reads (see Supp. Figure 7 for an example). We also identified a number of false positive variants called using the Minimap2 alignments that were corrected by variant calls using DuploMap alignments (see Supp. Figure 8 for an example).

**Figure 5:**
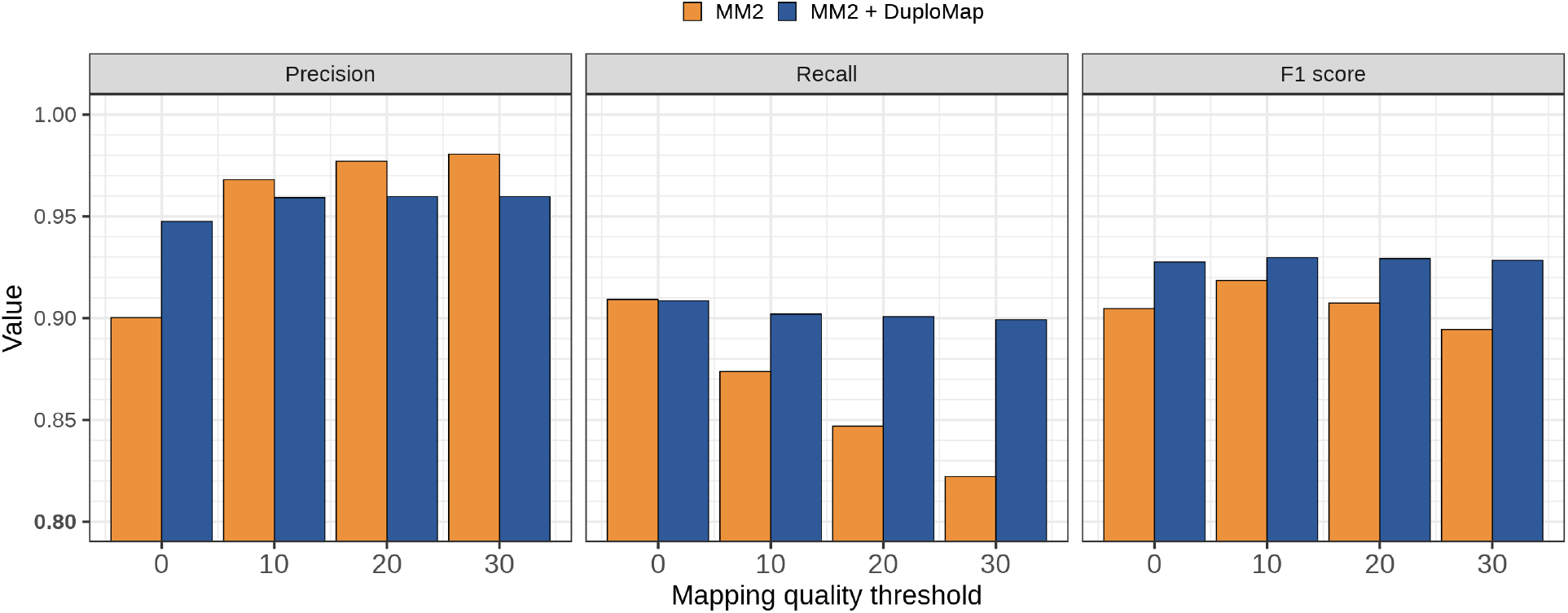
Comparison of variant calling accuracy for HG002 CCS reads using Minimap2 and Minimap2+DuploMap. SNVs were called using Longshot for different mapping quality thresholds. Precision, recall and *F*_1_ values were calculated by comparison to the GIAB v4.1 benchmark calls in the *Long-SegDups* regions that overlapped with the GIAB high-confidence regions.

Next, we directly compared the SNVs calls made on the HG002 CCS dataset using Minimap2 aligned reads and reads re-aligned using DuploMap. We utilized 10X Genomics linked-read variant calls for the same individual as an independent source for comparison. In *Long-SegDups* regions on chromosomes 1-22, 83,648 DuploMap-derived SNVs were shared with 10X calls compared to 72,830 for Minimap2. 14,713 SNVs called exclusively using the DuploMap alignments were supported by 10X calls. We also identified 211 calls that were shared between the DuploMap calls and 10X calls but were absent in the GIAB v4.1 benchmark calls. Visual inspection of these calls suggested that for many of them, the GIAB benchmark callset is either missing a variant or has the incorrect genotype (see Supp. Figure 9 for an example). In addition, 36,021 SNVs were located outside the GIAB v4.1 regions and were shared between all three callsets: 10X, Minimap2 and DuploMap. These variants are likely to be true positives that are located outside current GIAB high-confidence regions.

### Uninformative PSVs and variant calling using short reads

In addition to aligning reads that overlap segmental duplications, DuploMap also estimates genotypes for PSVs to identify unreliable or uniformative PSVs. Uninformative PSVs are likely to be the result of true variants in segmental duplications and therefore, should also be called as variants using long-read variant calling. Analysis of Longshot variant calls for the HG002 CCS dataset showed that 42.5% of the SNVs called using the DuploMap alignments in *Long-SegDups* regions intersected PSV sites such that the variant allele matched the PSV allele at the homologous site. Such non-reference PSVs were not specific to DuploMap alignments; 43.4% of the SNVs called using Minimap2 alignments also intersected PSVs. In both cases, approximately 76% of the variants overlapping PSVs are present in the dbSNP database (build 151) [36].

Next, to assess the impact of uninformative PSVs on short read variant calling, we analyzed PacBio CCS read data for the HG001 (NA12878) genome for which pedigree-derived variant calls have been generated by the Platinum Genomes (PG) Project using whole-genome Illumina sequence data [37]. We focused our analysis of PSVs on two-copy segmental duplications. Of the 14,800 PSVs in two-copy duplications - with high confidence genotypes (QUAL *≥* 60) estimated by DuploMap - 16.5% had a genotype of (0/0, 0/1) and 6.0% had a genotype of (0/0, 1/1). A genotype of (0/0, 1/1) for a PSV implies that the genomic sequence at both homologous positions (on both alleles) is identical and hence the PSV cannot differentiate between reads from the two homologous sequences. Such PSVs are expected to cause incorrect read mapping and lead to incorrect variant calls since short read mapping tools rely on PSVs to place reads with high confidence in segmental duplications. For example, if a true variant is present in the region flanking the PSV position with a non-reference genotype, short reads covering the variant and the PSV can be mismapped to the homologous location resulting in a false variant call (see Supp. Figure 10 for an illustration).

To search for false variants resulting from uninformative PSVs, we identified variants in the Platinum Genomes variant calls [37] for HG001 that were located near PSVs. Of the 2,769 variants that were located near uninformative PSVs in the PG calls, we identified 76 variants such that the variant was missing in the CCS variant calls but another variant was present at the homologous position with the same alternate allele. One such example of a false variant due to a uninformative PSV was located at the *PMS2* locus (Figure 6). The short-read PG calls report a SNV at chr7:6,752,118 (hg38 reference genome, rs1060836) that was also reported in gnomAD database of human variants [38] with an average allele frequency of 0.16 but with 7-fold lower homozygotes than expected – indicative of a false variant. This SNV was absent from long read variant calls but a SNV located at chr7:5,972,674 – the position homologous to chr7:6,752,118 – was present in the DuploMap based calls and also in the 10X Genomics variant calls. This SNV was located less than 75 bases from an uninformative PSV chr7:5,972,749 that was actually called as a variant with the variant allele being the same as the allele at the homologous site (Figure 6).

**Figure 6:**
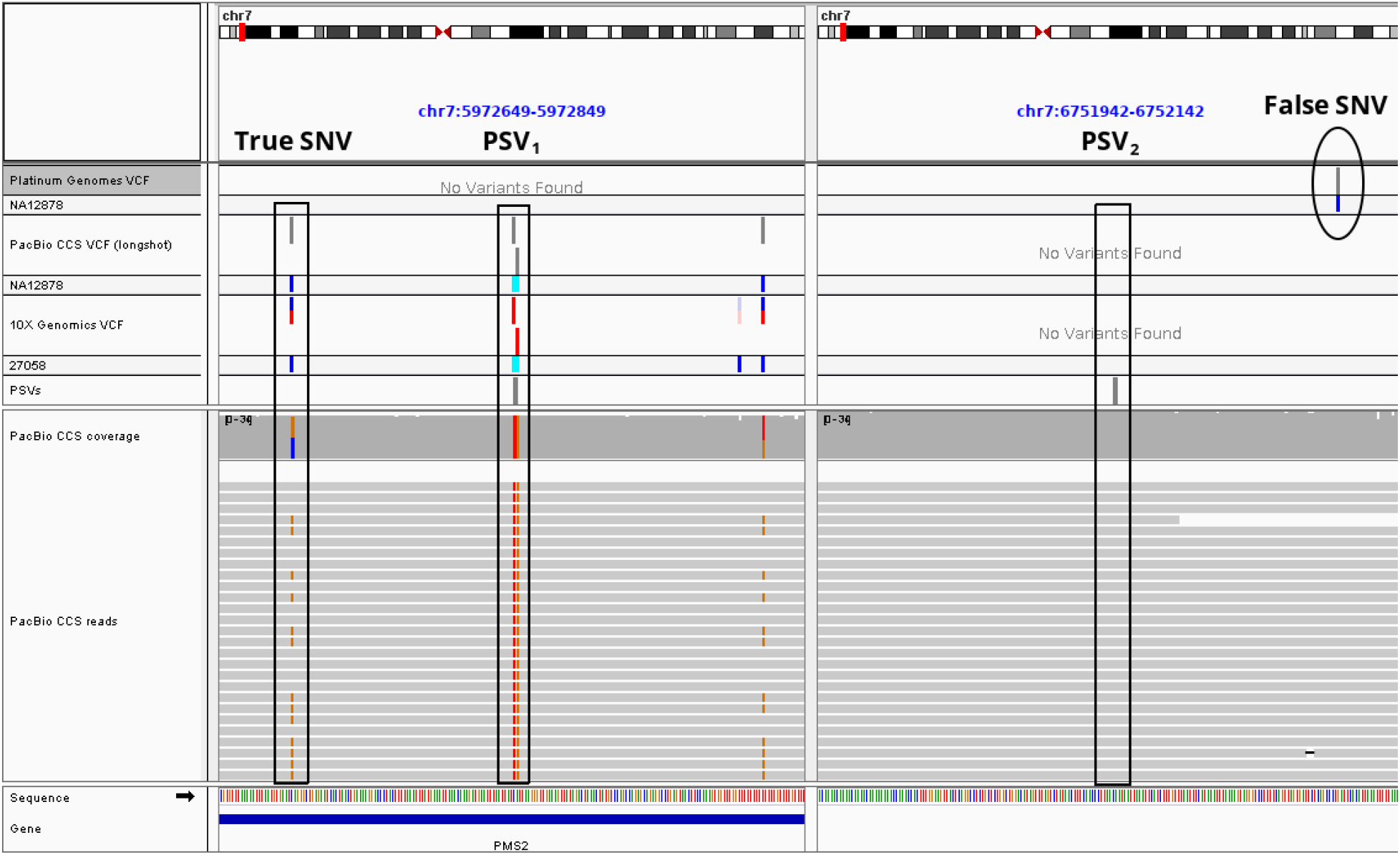
Illustration of how unreliable PSVs adversely impact short-read variant calling in segmental duplications. An Integrated Genomics Viewer (IGV) view of a duplicated region on chromosome 7 that overlaps the *PMS2* gene is shown. A PSV is located at chr7:5972749 (allele = with the homologous position at chr7:6752042. Variant calling on PacBio CCS reads (aligned with DuploMap) identifies two variants, a homozygous variant at the PSV site (chr7:5972749:CA:TG) and a heterozygous SNV located nearby (chr7:5972674:C:G). Both of these variants are supported by 10X Genomics variant calls but are absent from short-read variant calls for the same individual (Platinum Genomes VCF track). In addition, short-read variant calling results in a false SNV (chr7:6752118:G:C) at the position homologous to chr7:5972674 - a result of short read mismapping due to the unreliable PSV.

## Discussion

In this paper, we presented DuploMap, a method designed specifically for re-aligning SMS reads that are mapped to segmental duplications by existing long-read alignment tools in order to improve accuracy. A unique feature of DuploMap is that it jointly analyzes reads overlapping segmental duplications and explicitly leverages paralogous sequence variants or PSVs for mapping. Using whole-genome human data generated using multiple SMS technologies, we demonstrate that DuploMap significantly improves the mappability of reads overlapping long segmental duplications in the human genome. DuploMap is not a stand-alone long read alignment tool but complements existing tools such as Minimap2 that tend to be conservative in aligning reads in segmental duplications.

The development of DuploMap is motivated by the goal of using long read sequencing technologies for variant calling in long segmental duplications that are problematic for short-read sequencing. The Genome in a Bottle Consortium has developed high-confidence small variant call sets for reference human genomes [39, 33, 34]. Their first call sets were based on short read sequencing and hence exclude almost all segmental duplications. The GIAB consortium is expanding the small variant calls to repeats including segmental duplications using the PacBio CCS and 10X Genomics linked read data-types. Accurate and sensitive read mapping of long reads is a pre-requisite for accurate and sensitive variant calling in long repeats in the human genome. Variant calling using the DuploMap aligned reads identified 14,713 variants in segmental duplications that were shared with 10X Genomics variant calls but were not called using Minimap2 aligned reads. This indicates that DuploMap can prove useful for variant calling in segmental duplications using PacBio CCS reads.

DuploMap is a robust method that works for multiple long read sequencing technologies (PacBio and ONT), can handle reads with high and low error rates, and can post-process reads aligned with different long-read alignment tools. DuploMap’s approach of jointly modeling PSV genotypes and read alignments can potentially be used to improve the mapping of linked-reads in segmental duplications [40, 41, 42]. Although we have focused on variant calling, the ability to map long reads to segmental duplications with high sensitivity can benefit other uses of long read sequencing. Oxford Nanopore sequencing enables the detection of DNA methylation directly from the raw base signal [43, 44]. Miga et al. [25] have used a unique *k*-mer based mapping strategy to improve read mapping to generate base-level DNA methylation maps for the centromere of the X chromosome. DuploMap based alignment can enable the analysis of the methylation levels of duplicated genes that cannot be measured using short-read based methylation assays.

Analysis of PacBio CCS reads for a human genome showed that a significant number of PSVs overlap with variants and hence are uninformative for read mapping in segmental duplications. PSVs are defined based on the reference human genome sequence and a common variant in the human population can be incorrectly considered as a PSV if the variant allele is represented in the reference. In addition, gene conversion is well known to result in overlap between PSVs and variants [45, 46]. We also demonstrated that uninformative PSVs can cause incorrect mapping of short reads to homologous sequences resulting in both false positive and false negative variant calls. This problem can be alleviated by using information about reliable PSVs derived from analysis of long read datasets to inform short read mapping and variant calling in segmental duplications.

DuploMap has several limitations. First, the memory usage for DuploMap scales linearly with increasing number of reads since it stores information about all reads that overlap a single cluster of duplications. This can be reduced by writing some of the mapping information to disk or limiting the re-alignment to segmental duplications with low copy number. Second, DuploMap is not a stand-alone aligner and starts from alignments provided by existing long-read alignment tools. If a read is not aligned or aligned to a location that is not homologous to its correct location, DuploMap cannot find the correct alignment. Third, DuploMap does not currently account for missing sequences or copy number changes. Segmental duplications are well known to be hotspots of copy number variation and large structural variants in the human genome [26, 47]. Copy number information about duplicated sequences can be estimated using tools such as Quic*k*-mer2 [48] and used to potentially improve long read mapping in segmental duplications. Finally, DuploMap relies on segmental duplications identified from a reference genome. The current human reference assembly has many gaps and unassembled repetitive sequences. Ongoing efforts to generate gapless assemblies of human chromosomes using a combination of sequencing technologies [25] are expected to generate a more complete reference human genome sequence which in turn will enable more accurate long read alignment in segmental duplications.

## Competing interest statement

The authors declare no competing interests.

## Data availability

All datasets analyzed in this paper have been generated previously and are publicly available (links provided in Supplementary Data). DuploMap is implemented in the Rust programming language and is freely available for download at https://gitlab.com/tprodanov/duplomap. DuploMap can be used to map reads in individual clusters of segmental duplications or across the entire genome. It is also available via conda (conda install -c tprodanov duplomap). The repository also contains links to pre-computed PSV databases and BED files with *Long-SegDups* for the hg19 and hg38 versions of the human genome.

## Acknowledgements

The research was supported by the National Human Genome Research Institute of the National Institute of Health under award number R01HG010149. The content is solely the responsibility of the authors and does not necessarily represent the official views of the National Institutes of Health. We thank Justin Zook and Justin Wagner for useful discussions and help with accessing GIAB datasets.

## Notes

### Competing Interest Statement

The authors have declared no competing interest.

