## Supplementary Data and Methods for "Sensitive alignment using paralogous sequence variants improves long read mapping and variant calling in segmental duplications"

### Supplementary Figures and Tables

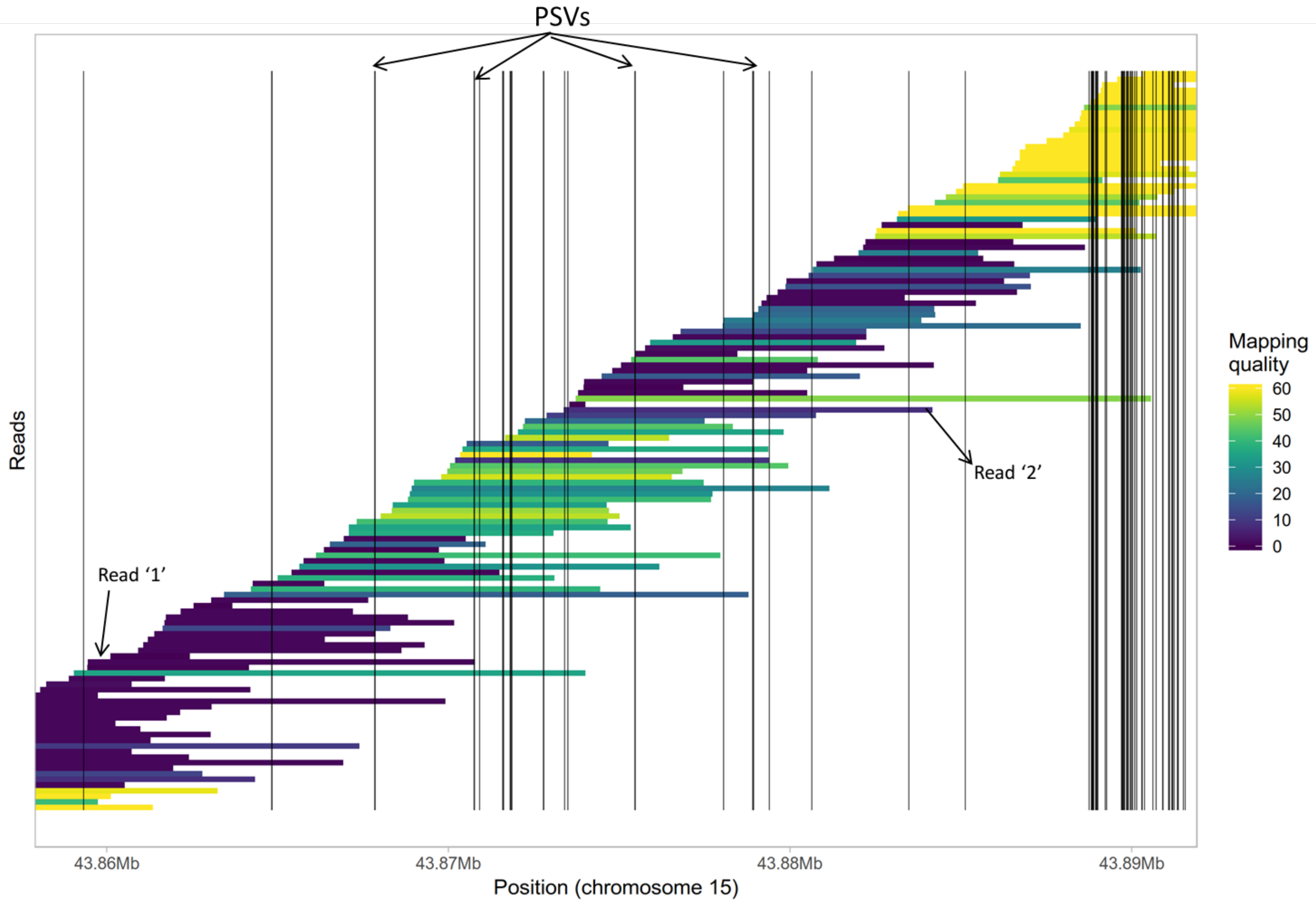


**Supplementary Figure 1: Illustration of sub-optimal mapping of long reads in segmental duplications.** Reads mapped using the Minimap2 aligner to a 35 kb region from a segmental duplication on chromosome 15 (covering the STRC gene) are shown. Reads are shown as horizontal bars (color-coded by mapping quality) while PSVs are shown as vertical lines. Several reads overlap multiple PSVs (e.g. read '2' overlaps 6 PSVs) but are still assigned low mapping quality. Other reads overlap no PSVs (e.g. read '1') and hence cannot be mapped uniquely.

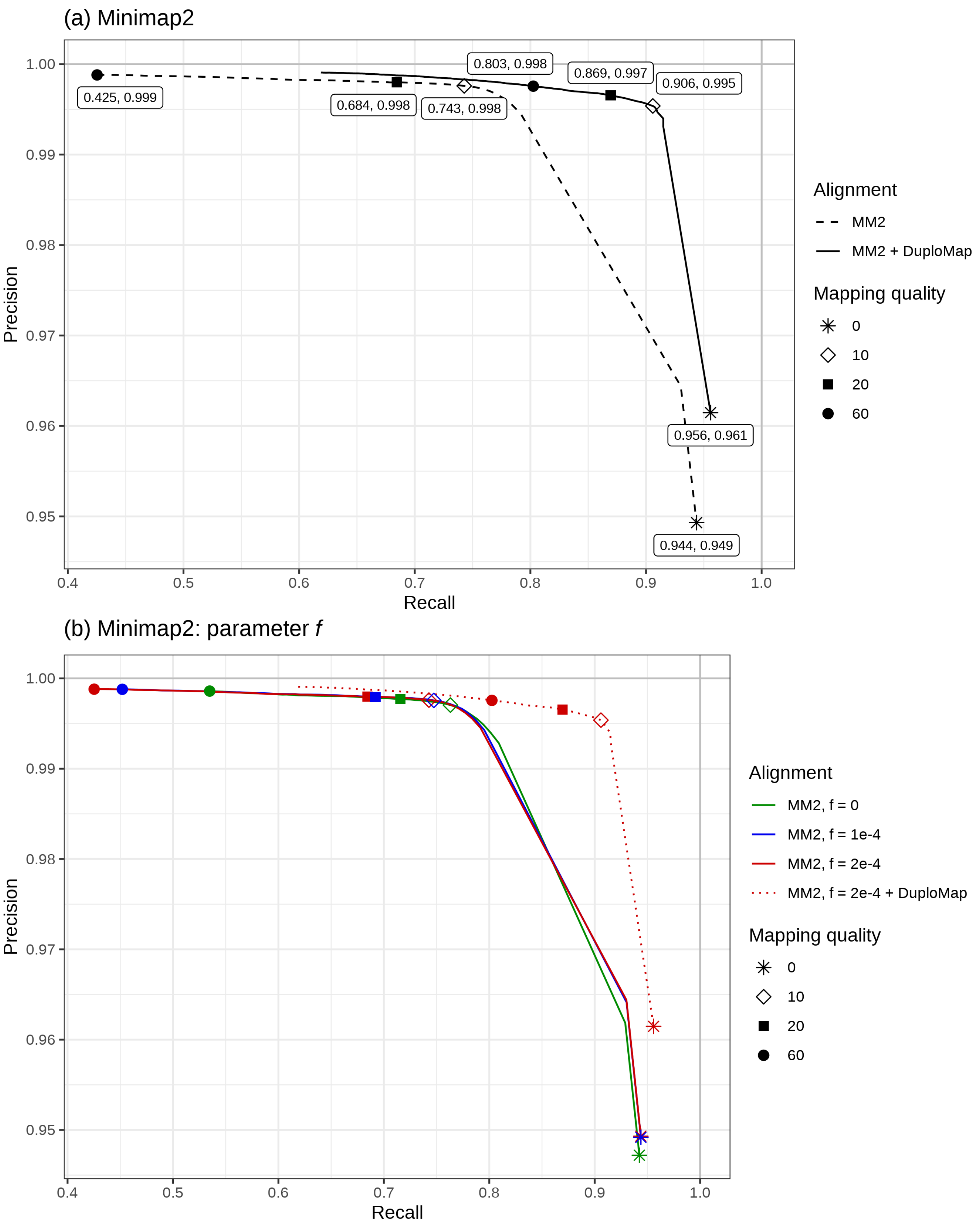


**Supplementary Figure 2: Accuracy of read mapping using Minimap2 and Minimap2+DuploMap on simulated long read data in segmental duplications.** Reads of median length 8.5 kb were used for simulations. (a) Accuracy of Minimap2 and Minimap2 + DuploMap alignments. (b) MM2 accuracy with different values of parameter $f$ (discarding top $f$ of the repetitive minimizers, $2 \cdot 10^{-4}$ by default).

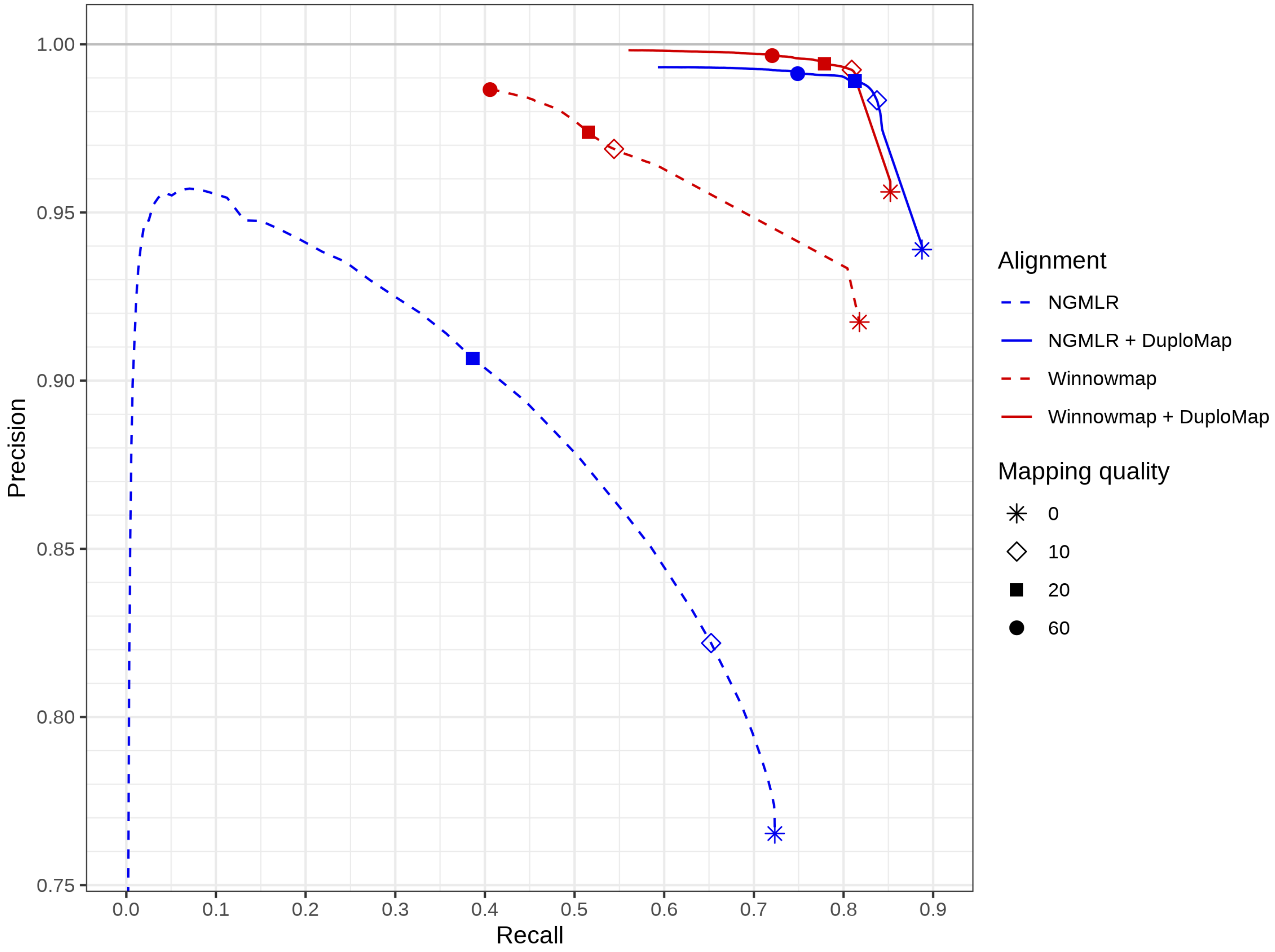


**Supplementary Figure 3: Improvement in accuracy of read mapping in segmental duplications using DuploMap in combination with the NGMLR and Winnowmap alignment tools.** Each precision-recall curve is plotted using different mapping quality thresholds.

| Length (kb) | Aligner | Running time (hh:mm) | Mapping speed (reads per second) | Memory usage (Gb) |
|---|---|---|---|---|
| 8.5 | Minimap2 | 3:54 | 105.1 | 13.33 |
| 8.5 | NGMLR | 35:18 | 11.2 | 10.55 |
| 8.5 | BLASR | - | 3.6 | 29.29 |
| 8.5 | Minimap2, $f = 0$ | - | 15.6 | 67.68 |
| 8.5 | DuploMap | 1:39 | 10.7 | 20.16 |
| 20 | Minimap2 | 3:57 | 44.9 | 15.11 |
| 20 | DuploMap | 1:27 | 4.9 | 17.58 |
| 50 | Minimap2 | 2:57 | 19.4 | 15.38 |
| 50 | DuploMap | 1:30 | 2.1 | 16.95 |

**Supplementary Table 1: Running time and memory usage of long read alignment tools and DuploMap on simulated SMS reads.** Running time shows the elapsed real time for each aligner using 8 cores. Mapping speed shows the average number of reads analyzed per second (by a single core). Note that DuploMapanalyses only a subset of reads that intersect segmental duplications. Only the subset of reads that intersect segmental duplications were mapped using BLASR and Minimap2 with $f = 0$ due to their long running time. All tools were run on a CentOS 6.6 system with Intel Xeon CPU E5-2670 @ 2.60 GHz, with jobs managed by a Torque/PBS system.

| Genome | Sequencing technology | Median coverage | Running time (hh:mm) | Memory usage (Gb) |
|---|---|---|---|---|
| HG001 | ONT | 36 | 4:57 | 38.29 |
| HG001 | PacBio CCS | 29 | 4:09 | 29.54 |
| HG002 | PacBio CLR | 45 | 8:31 | 61.88 |
| HG002 | PacBio CCS | 29 | 6:29 | 25.79 |
| HG002 | ONT | 58 | 14:28 | 55.91 |
| HG003 | PacBio CLR | 20 | 3:54 | 32.35 |
| HG004 | PacBio CLR | 19 | 3:24 | 29.13 |
| HG005 | PacBio CCS | 32 | 5:51 | 30.58 |

**Supplementary Table 2: Running time and memory usage of DuploMap on real data.** Running time represents elapsed real time using 8 cores.

| Non-ref PSVs (%) | PSV positions genotyped (%) | Incorrect genotypes (%) | Precision (%) | Recall (%) |
|---|---|---|---|---|
| 0 | 86.6 | 0.000 | 99.91 | 86.85 |
| 15 | 85.0 | 0.001 | 99.90 | 86.54 |
| 30 | 81.3 | 0.000 | 99.91 | 86.17 |

**Supplementary Table 3: Simulations with non-reference PSVs.** Two-copy segmental duplications in the human genome (hg19) were used for assessing the impact of unreliable PSVs (non-reference) on the accuracy of DuploMap. Of the 52,276 high-complexity PSVs in two-copy segmental duplications, we modified the genome sequence for one of the two copies for 0, 15 and 30% of the PSVs. Reads were simulated using the modified genome and mapped using DuploMap. The percentage of PSV positions (total count = 104,552) with high quality genotypes (all filters pass and quality score $\geq 60$) is shown in column 2. The precision and recall for reads mapped with mapping quality $\geq 30$ is also shown.

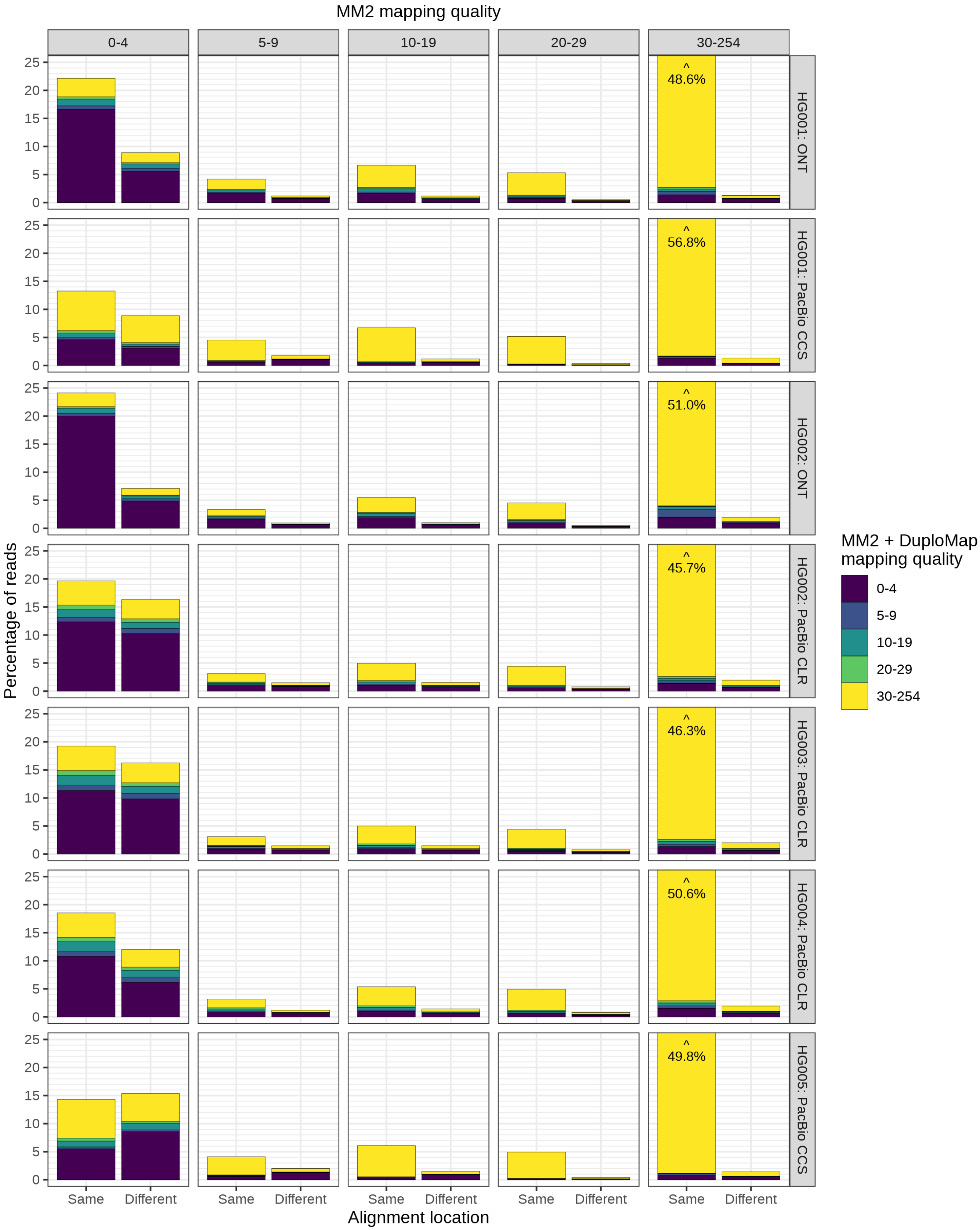


**Supplementary Figure 4: Comparison of mapping qualities and alignment locations for reads aligned with Minimap2 and Minimap2 + DuploMap on multiple long-read datasets.** Column contain reads with corresponding mapping quality in the MM2 alignments. Two bars in each subplot represent reads that have same or different alignments in MM2 and MM2 + D. Bar height represents percentage of reads in the corresponding category out of all analyzed reads in the dataset, and color shows alignment mapping quality after MM2 + D. Some bars are clipped, in that cases total bar height is shown at the top of the bar.

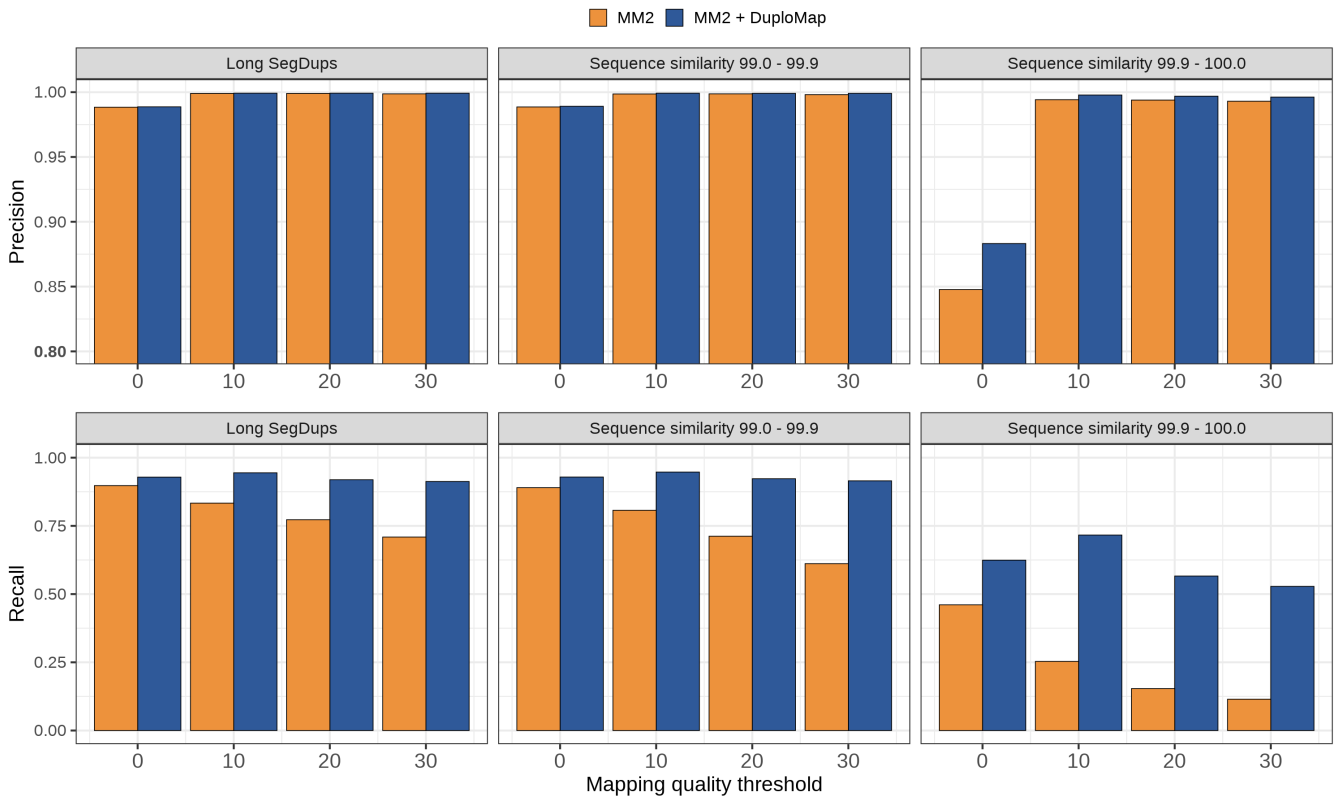


**Supplementary Figure 5: Precision and recall of variant calling in segmental duplications using simulated reads aligned with Minimap2 and Minimap2 + DuploMap.** Three columns show different subsets of variants: within all *Long-SegDups* regions; within *Long-SegDups* regions with sequence similarity between 99.0% and 99.9%; and within *Long-SegDups* regions with sequence similarity between 99.9% and 100%.

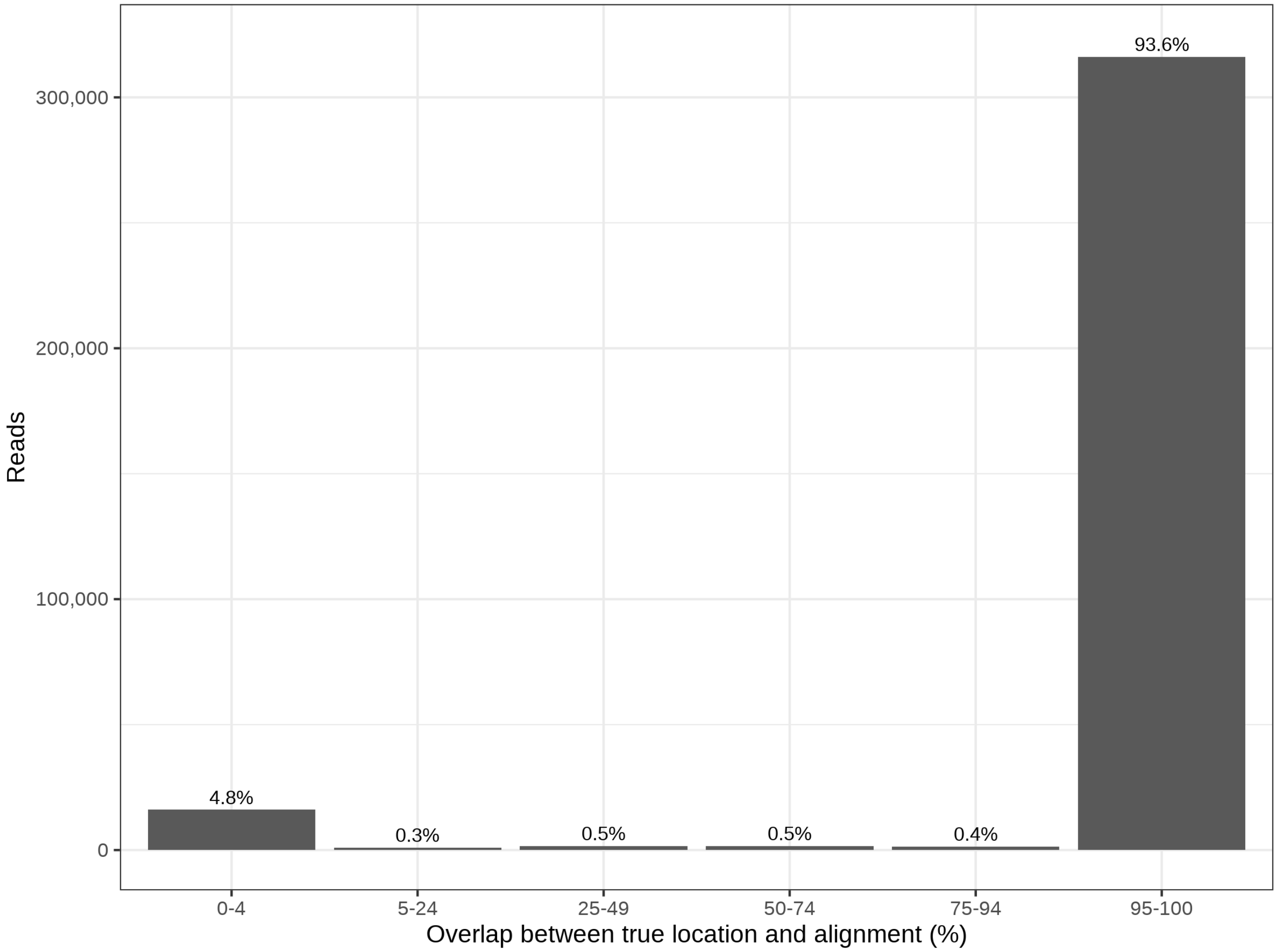


**Supplementary Figure 6: Overlap percentage between true location and alignment locations in Minimap2 mapping of long simulated reads.**

| Genome | Alignment method | Genome subset | Variant quality | Number of variants | Precision | Recall | F1 |
|---|---|---|---|---|---|---|---|
| HG002 | MM2 | v3 | 30 | 3,010,414 | 0.9963 | 0.9900 | 0.9931 |
| | MM2 + DuploMap | v3 | 30 | 3,010,534 | 0.9963 | 0.9900 | 0.9932 |
| | MM2 | v4 | 30 | 3,319,220 | 0.9973 | 0.9837 | 0.9904 |
| | MM2 + DuploMap | v4 | 30 | 3,320,654 | 0.9972 | 0.9840 | 0.9905 |
| | MM2 | v4 ∩ SD | 30 | 36,044 | 0.9680 | 0.8738 | 0.9185 |
| | MM2 + DuploMap | v4 ∩ SD | 30 | 37,548 | 0.9592 | 0.9020 | 0.9297 |
| | MM2 | v4 ∩ SD | 60 | 35,902 | 0.9701 | 0.8723 | 0.9186 |
| | MM2 + DuploMap | v4 ∩ SD | 60 | 37,421 | 0.9611 | 0.9007 | 0.9299 |
| | MM2 | v4 ∩ SD | 90 | 35,582 | 0.9727 | 0.8668 | 0.9167 |
| | MM2 + DuploMap | v4 ∩ SD | 90 | 37,192 | 0.9634 | 0.8974 | 0.9292 |

**Supplementary Table 4: Comparison of variant calling accuracy for HG002 CCS reads aligned with Minimap2 (MM2) and DuploMap.** SNVs were called using Longshot (mapping quality threshold of 10). 'v3' refers to the high-confidence regions of the genome in the GIAB v3.3.2 call set for each genome and 'v4' refers to the expanded high-confidence regions in the GIAB v4.1 callset for HG002. 'SD' or *Long-SegDups* refers to the genomic regions in which reads were realigned using DuploMap.

| Gene | Chromosome | Sum exon length | MM2 coverage (%) MQ $\geq$ 10 | MM2 coverage (%) MQ $\geq$ 20 | $\Delta$ MM2 + D coverage (%) MQ $\geq$ 10 | $\Delta$ MM2 + D coverage (%) MQ $\geq$ 20 |
|---|---|---|---|---|---|---|
| NAIP | 5 | 7,704 | 20.6 | 9.7 | +79.4 | +90.3 |
| C4B | 6 | 5,427 | 36.8 | 28.6 | +63.2 | +71.4 |
| SMN1 | 5 | 2,234 | 0.0 | 0.0 | +59.1 | +59.1 |
| GTF2I | 7 | 5,889 | 55.0 | 52.6 | +45.0 | +47.4 |
| C4A | 6 | 5,427 | 57.9 | 53.6 | +42.1 | +46.4 |
| GTF2IRD2 | 7 | 5,394 | 48.3 | 22.0 | +18.7 | +45.0 |
| PPIP5K1 | 15 | 6,575 | 90.3 | 81.6 | +9.7 | +18.4 |
| CATSPER2 | 15 | 4,538 | 95.2 | 95.2 | +4.8 | +4.8 |
| PDPK1 | 16 | 8,106 | 95.3 | 93.7 | +4.7 | +6.3 |
| SMN2 | 5 | 2,671 | 62.9 | 57.1 | +4.5 | +10.3 |
| NEB | 2 | 26,310 | 99.5 | 98.0 | +0.5 | +2.0 |
| OTOA | 16 | 4,180 | 100.0 | 96.3 | +0.0 | +3.7 |
| CFC1 | 2 | 1,669 | 100.0 | 0.0 | +0.0 | +100.0 |
| OCLN | 5 | 6,549 | 100.0 | 94.1 | +0.0 | +5.9 |
| PMS2 | 7 | 5,150 | 97.9 | 85.4 | +0.0 | +12.5 |
| NCF1 | 7 | 2,022 | 100.0 | 93.0 | +0.0 | +7.0 |
| CR1 | 1 | 9,953 | 86.4 | 85.4 | -1.0 | +0.0 |

**Supplementary Table 5: Mappability of 17 disease-associated genes with Minimap2 and Minimap2 + DuploMap for HG002 PacBio CCS dataset.** MM2 coverage columns show percentage of bases covered by at least 15 reads (half the median coverage) with high mapping quality ($\geq$ 10 and $\geq$ 20) in Minimap2 alignments in all exons of the corresponding gene. Last two columns show difference between percentage of covered bases in *Minimap2 + DuploMap* and *Minimap2* alignments.

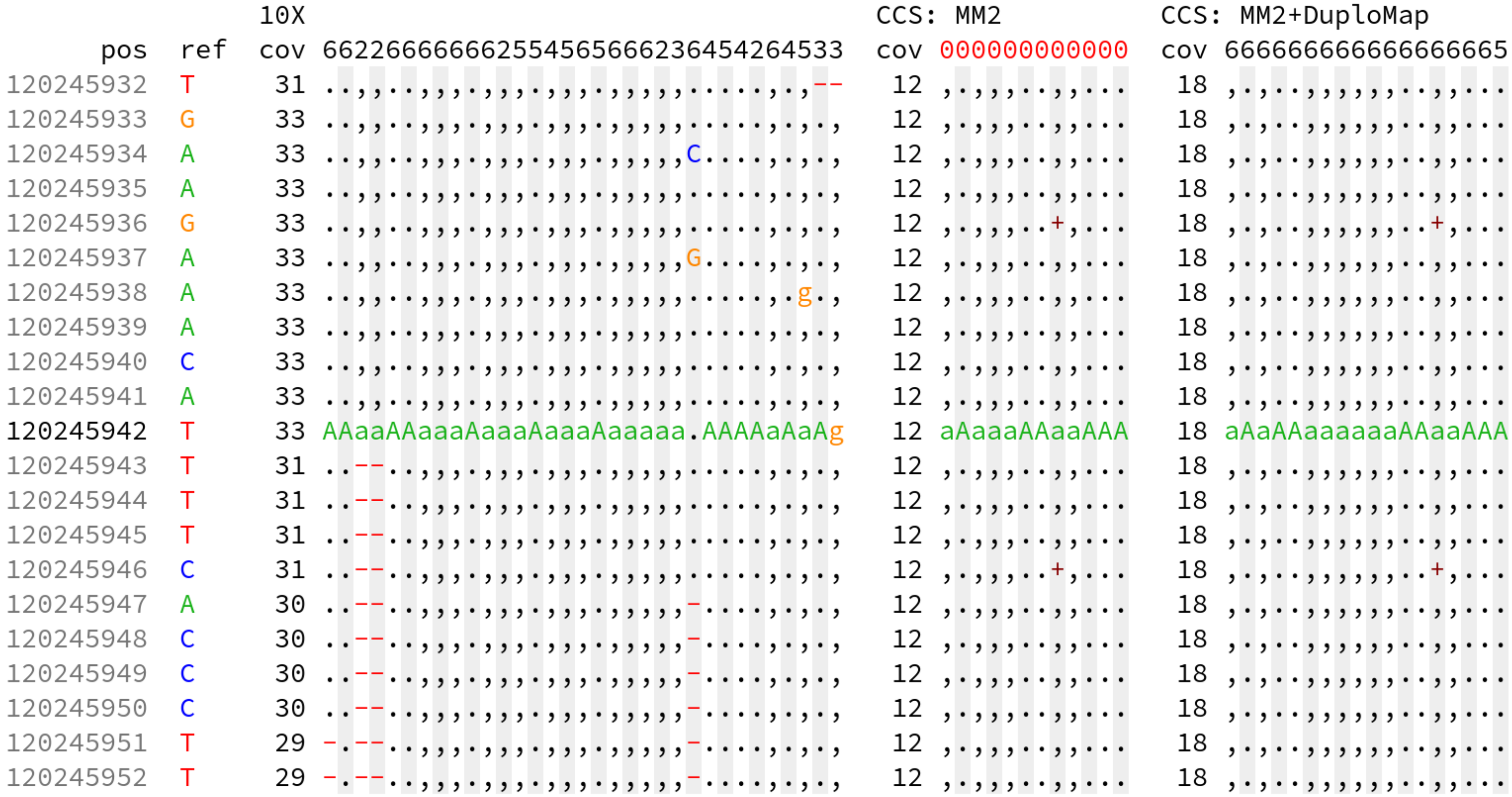


**Supplementary Figure 7: Example of a variant outside the GIAB high-confidence region identified using Minimap2 + DuploMap alignments of PacBio CCS reads.** Pileups of 10X Genomics linked-reads and PacBio CCS reads for the individual HG002 (aligned with Minimap2 and Minimap2 + DuploMap) in a window around the position chr1:120,245,942 (hg38) are shown. Each row shows a single position, and each column represents a single read. First digit of mapping quality is shown on top (0-6) and is highlighted in red for reads with mapping quality less than 10. The variant lies within 333kb duplication with sequence similarity 99.2%. The variant is present in the GIAB and 10X Genomics calls with genotype equal to 1/1. However, all CCS reads mapped using Minimap2 have low MAPQ and hence no variant is called. Minimap2 + DuploMap alignments have high mapping quality at this locus enabling Longshot to identify the variant with the correct genotype.

t

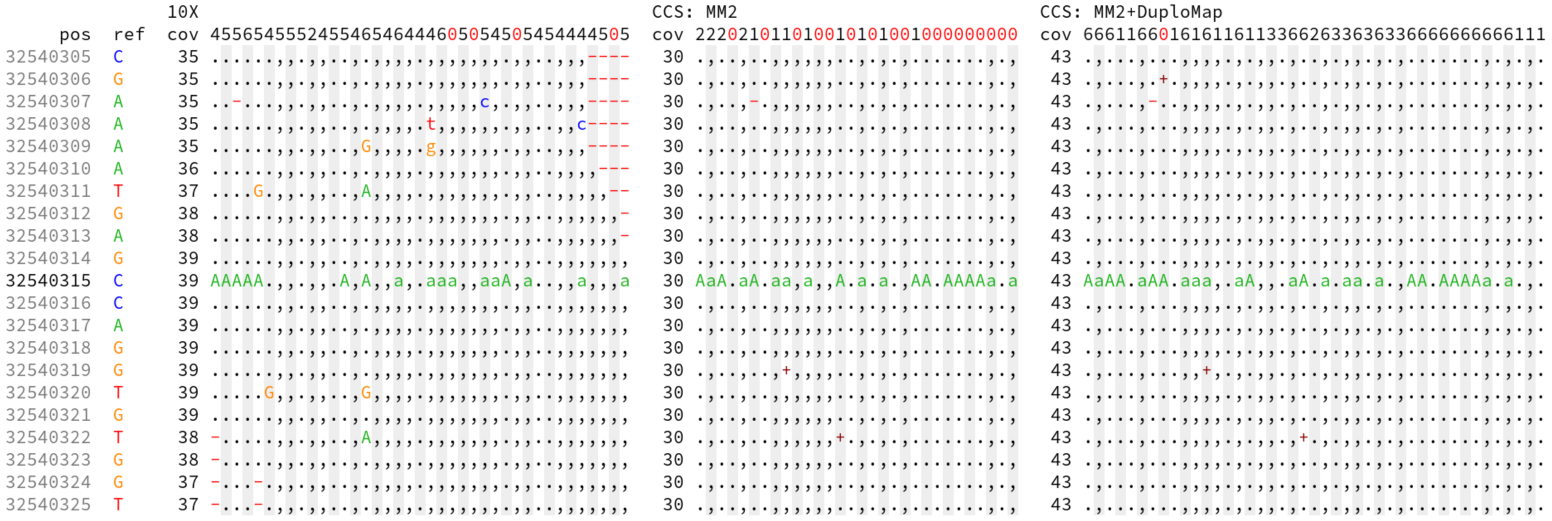


**Supplementary Figure 8: Example of how the improved mappability of DuploMap reduces false negatives in variant calling using PacBio CCS reads.** Pileups of 10X Genomics linked-reads and PacBio CCS reads for the individual HG002 (aligned with Minimap2 and Minimap2 + DuploMap) in a window around the position chr15:32,540,315 (hg38) are shown. Each row shows a single position, and each column represents a single read. First digit of mapping quality is shown on top (0-6) and is highlighted in red for reads with mapping quality less than 10. The variant lies within a 218kb duplication with sequence similarity 99.5%. The variant is present in the GIAB benchmark variant calls and the 10X Genomics calls with genotype equal to 0/1. However, all CCS reads mapped using Minimap2 that have high mapping quality have the alternative allele 'A' resulting in a homozygous variant call (genotype = 1/1). After realignment using DuploMap, all reads have high mapping quality and the variant is called using Longshot with the correct genotype (0/1).

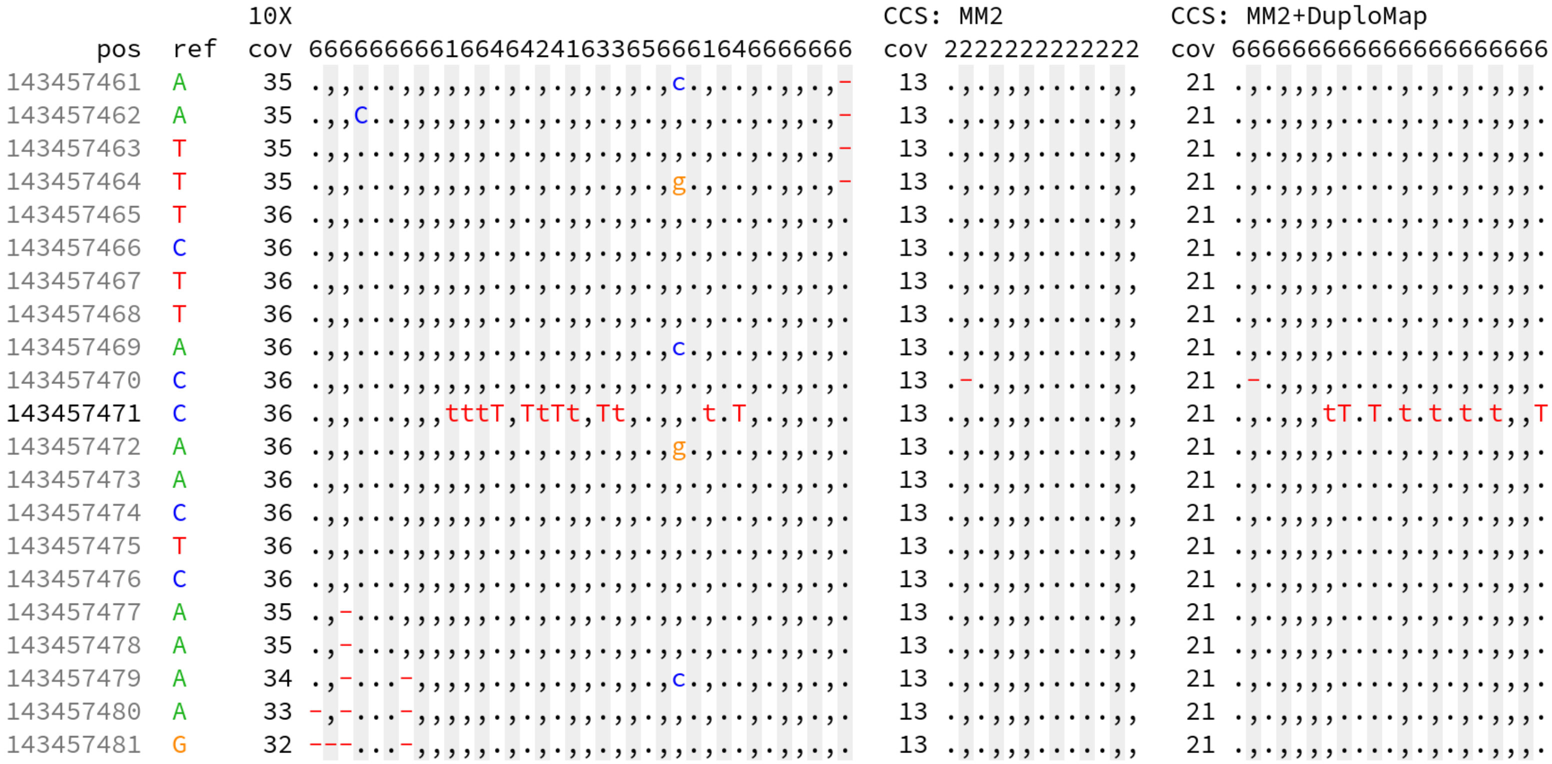


**Supplementary Figure 9: Example of a potential false negative call in the GIAB benchmark variant call set identified using DuploMap alignments.** Pileups of 10X Genomics linked-reads and PacBio CCS reads for the individual HG002 (aligned with Minimap2 and DuploMap) in a window around the position chr1:143,457,471 (hg38) are shown. Each row shows a single position, and each column represents a single read. First digit of mapping quality is shown on top (0-6). The position lies within a 220 kb long segmental duplication with sequence similarity 99.6%. The variant lies in high-confidence GIAB regions, but is absent in the GIAB benchmark variant calls. Nevertheless, the variantt is present in the 10X Genomics calls. However, all CCS reads mapped using Minimap2 have reference allele 'C' and hence the variant is not called. After realignment using DuploMap, the number of reads covering this position increases from 13 to 21 and includes 8 reads that support alternative allele 'T'. Using Longshot, a variant is called at this position that matches the 10X variant call.

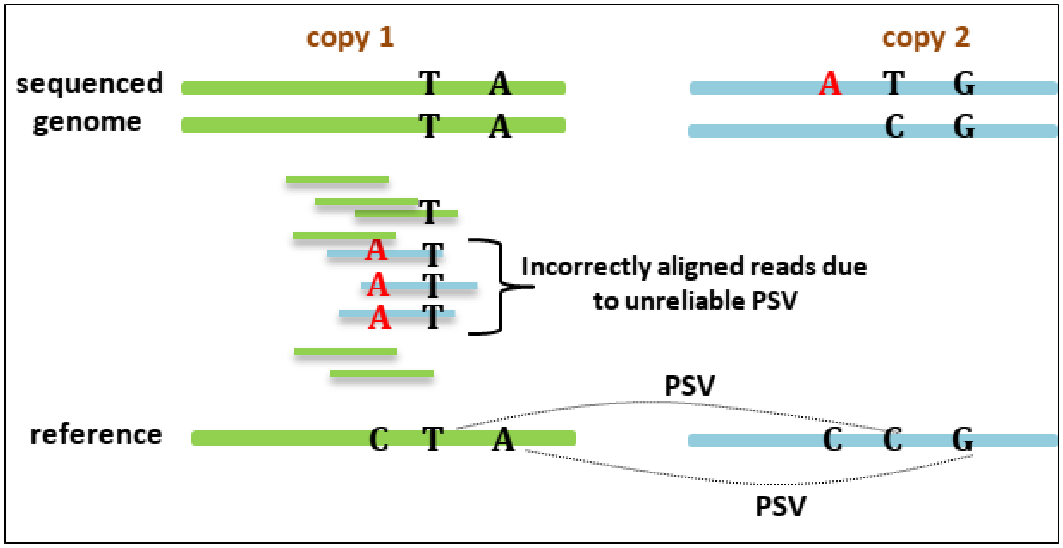


**Supplementary Figure 10: Illustration of how incorrect short read mapping due to unreliable PSVs can lead to false positive and false negative variant calls**. A two-copy segmental duplication is shown with two PSVs that distinguish ‘copy 1’ and ‘copy 2’. The sequenced genome carries a variant (A allele) on one of the haplotypes of ‘copy 2’. One of the two PSVs is actually a variant in the sequenced genome with the ’T’ allele instead of the ’C’ allele in ‘copy 2’. Hence, reads with the A allele that originate from ‘copy 2’ are mismapped to ‘copy ‘1’ resulting in a false positive variant call at the homologous position in ‘copy 1’ with the alternate allele being identical to the PSV allele.

### Supplementary Methods

#### Filtering PSVs

To identify low-complexity PSVs we count the number of unique $k$-mers (with $k = 3$) in a window around the PSV. PSVs for which the number of $k$-mers divided by the maximal number of $k$-mers for the window of the same size is less than 60% for substitutions and 80% for indels are filtered out. We also filter out PSVs for which it is difficult to distinguish between the two alleles due to high sequencing error rates. For a read $r$ that covers a PSV $v$, we calculate the alignment probabilities for each of the two alleles of the PSV $s_v^{(i)} = P(r_v \mid S_v^{(i)})$ and $s_v^{(j)} = P(r_v \mid S_v^{(i)})$. We say that the read has an *ambiguous* alignment for the PSV if $\max\{s_v^{(i)}, s_v^{(j)}\} / \min\{s_v^{(i)}, s_v^{(j)}\} < 4$. After the first iteration of the DuploMap algorithm, we remove all PSVs for which 30% or more reads have an ambiguous alignment. This filtering removes PSVs that were not identified as low-complexity but still have noisy local realignment probabilities. It is possible that the PSV in the sequenced genome has a sequence different from the two known alleles $S_v^{(i)}$ and $S_v^{(j)}$. This step can also filter out such PSVs.

#### Identifying candidate alignment locations for a read

In the PSV database, we store each pair of homologous sequences as a collection of pairs of windows $(w^{(1)},\ w^{(2)})$, where each window is approximately 100 bp in length. The windows are constructed from the pairwise alignment such that window $w^{(1)}$ in one of the sequences is aligned to the window $w^{(2)}$ in the other. For an aligned read $r$, we consider windows $\left\{w_i^{(1)}\right\}_{i=1}^{n}$ that intersect its primary alignment. Using the database, we can identify all windows $\left\{w_i^{(2)}\right\}_{i=1}^{n}$ that are homologous to the windows of the primary alignment of the read. Without loss of generality, suppose that all pairs of windows are on same strand in the genome. We reorder the indices so that windows $\left\{w_i^{(2)}\right\}_{i=1}^{n}$ are sorted by their genomic positions. Additionally, we define a function $pos_1\left(w^{(2)}\right)$ that returns a genomic position of the window $w^{(1)}$. To identify possible alignment locations we search for pairs of indices $i \le j$ such that

1. windows $w_i^{(2)}$ and $w_j^{(2)}$ have the same order in the read: $pos_1\left(w_i^{(2)}\right) \le pos_1\left(w_j^{(2)}\right)$,
2. location is not too short: $j \ge i + m$, where $m$ is half of the number of non-overlapping windows in the initial alignment,
3. location generated from windows $w_i^{(2)}$ and $w_j^{(2)}$ is not more than 20% longer than the biggest of the read length and the initial alignment size,
4. no other pair of indices $i' \le i,\ j' \ge j$ produces a possible alignment location.

For an existing primary alignment with start $x_l$, end $x_r$ and soft clipping $y_l$ and $y_r$ we generate an alignment location by adding padding of size $\max\left\{0, y_l + pos_1\left(w_i^{(2)}\right) - x_l\right\}$ to the left of the window $w_i^{(2)}$. Similarly, we add padding of size $\max\left\{0, y_r + x_r - pos_1\left(w_j^{(2)}\right)\right\}$ to the right of the window $w_j^{(2)}$.

#### LCS-based filtering of alignment locations

We filter possible alignment locations using longest common subsequences (LCS) between the $k$-mers of the read and the $k$-mers of each candidate alignment location ($k = 11$, by default). The LCSk++ algorithm [1] is used to find the LCS. If one or more of the alignment locations for a read is located near a gap or missing sequence in the reference genome, the LCS score may not reflect the alignment of the full sequence of the read. To avoid this behavior, we compute the LCS scores for a pair of locations using a truncated read sequence. The read is truncated using the location of the last (or first) $k$-mer that is shared between the read and both locations.

To select a smallest non-empty subset of alignment locations that dominate all other locations we construct a directed graph, where each node represents a single location. For a pair of locations $i$ and $j$ if location $i$ dominates location $j$ we add an edge from the node $j$ to node $i$ (worse to best). We add edges in both directions if neither location dominates. Afterwards, we split the graph on strongly connected components [2] and select all locations from the sink component.

#### Identifying reads with high discordance with PSVs

To find reads that show high discordance with PSVs, we calculate the number of conflicts (mismatches) between each read and the PSVs it intersects. For a given read $r$ mapped to location $i$, we analyse all reliable PSVs that intersect the new primary alignment location for the read. The second position for different PSVs may lie in different homologous locations (denoted by $(-i)$). We define the conflict rate for the read $r$ as

$$\frac{\sum_v \mathbb{1}\left(s_v^{(-i)}/s_v^{(i)} \geq 10\right)}{\sum_v \mathbb{1}\left(\max\{s_v^{(i)}, s_v^{(-i)}\}/\min\{s_v^{(i)}, s_v^{(-i)}\} \geq 10\right)},$$

where $\mathbb{1}$ denotes indicator function, $s_v^{(i)} = P(r_v \mid S_v^{(i)})$ and $s_v^{(-i)} = P(r_v \mid S_v^{(-i)})$ represent alignment probabilities for two alleles of the PSV $v$. In the above formula, the denominator represents the numbers of PSVs that have big difference between alignment probabilities for two alleles. The value in the numerator shows the number of PSVs that do not support location $i$.

For a given cluster of segmental duplications, we estimate the average conflict rate using all reads mapped to the cluster with high mapping quality and with at least five PSVs. We use the average conflict rate and the binomial test to test if the observed number of conflicting PSVs is higher than expected. Reads for which the Bonferroni-corrected p-value is lower than 0.05 are assigned a low mapping quality (5 by default).

#### Mappability of exons

To calculate the mappability of disease-associated genes using long reads, we calculated the percentage of positions covered by at least 10 reads with mapping quality greater than a specific threshold. We only analysed positions that were located in at least one exon of the GENCODE annotation for that gene [3].

#### Estimating coverage

To calculate read coverage for PacBio and Oxford Nanopore whole-genome sequencing we selected 200,000 positions at random for the hg38 genome (using `bedtools random`). We then selected 100,000 positions (at random) that lie on chromosomes 1-22 outside of centromeres and telomeres. For each position $x$ we counted the number of reads (passing samtools flag 3844) with alignment starting at position $\leq x$ and ending at position $\geq x$. Then, the median value of the measured coverages was taken.

#### Pileups

We constructed pileups using the pileuppy tool `v0.2.1` available at `https://gitlab.com/tprodanov/pileuppy`.

#### Datasets

Alignment and variant calling files can be found at the following links:

- HG002 (NA24385) PacBio CLR: https://ftp-trace.ncbi.nih.gov/giab/ftp/data/AshkenazimTrio/HG002_NA24385_son/PacBio_MtSinai_NIST/PacBio_minimap2_bam
- HG002 (NA24385) PacBio CCS: https://ftp-trace.ncbi.nih.gov/giab/ftp/data/AshkenazimTrio/HG002_NA24385_son/PacBio_CCS_15kb/GRCh38_no_alt_analysis
- HG002 (NA24385) Oxford Nanopore: https://ftp-trace.ncbi.nlm.nih.gov/giab/ftp/data/AshkenazimTrio/HG002_NA24385_son/Ultralong_OxfordNanopore/combined_2018-08-10
- HG002 (NA24385) 10X: https://ftp-trace.ncbi.nih.gov/giab/ftp/data/AshkenazimTrio/analysis/10XGenomics_ChromiumGenome_LongRanger2.2_Supernova2.0.1_04122018/GRCh38
- HG002 (NA24385) GIAB benchmark calls: https://ftp-trace.ncbi.nlm.nih.gov/giab/ftp/release/AshkenazimTrio/HG002_NA24385_son.
- HG003 (NA24149) PacBio CLR: https://ftp-trace.ncbi.nlm.nih.gov/giab/ftp/data/AshkenazimTrio/HG003_NA24149_father/PacBio_MtSinai_NIST/PacBio_minimap2_bam
- HG004 (NA24143) PacBio CLR: https://ftp-trace.ncbi.nlm.nih.gov/giab/ftp/data/AshkenazimTrio/HG004_NA24143_mother/PacBio_MtSinai_NIST/PacBio_minimap2_bam
- HG005 (NA24631) PacBio CCS: https://ftp-trace.ncbi.nih.gov/giab/ftp/data/ChineseTrio/HG005_NA24631_son/PacBio_SequelII_CCS_11kb/HG005_GRCh38
- HG005 (NA24631) GIAB benchmark calls: https://ftp-trace.ncbi.nlm.nih.gov/giab/ftp/release/ChineseTrio/HG005_NA24631_son
- HG001 (NA12878) Oxford Nanopore: https://github.com/nanopore-wgs-consortium/NA12878
- HG001 (NA12878) PacBio CCS: https://ftp-trace.ncbi.nih.gov/giab/ftp/data/NA12878/PacBio_SequelII_CCS_11kb
- HG001 (NA12878) 10X: https://ftp-trace.ncbi.nih.gov/giab/ftp/data/NA12878/10Xgenomics_ChromiumGenome_LongRanger2.1_09302016/NA12878_GRCh38
- HG001 (NA12878) Platinium Genome: ftp://ussd-ftp.illumina.com/2017-1.0/hg38/small_variants/NA12878/
